# Extracellular vesicles as indicators of environmental stress response in *Lactiplantibacillus plantarum*: a multi-platform study

**DOI:** 10.1101/2025.11.21.689665

**Authors:** Agnieszka Razim, Astrid Laimer-Digruber, Anna M. Schmid, Tanja V. Edelbacher, Magdalena E. Paschal, Tamara Weinmayer, Michael Thaler, Magdalena E. Skalska, Paweł Migdał, Mattia Morandi, Catherine Daniel, Dagmar Srutkova, Martin Schwarzer, Jiri Hrdy, Monika Ehling-Schulz, Sabina Górska, Aleksandra Inic-Kanada, Ursula Wiedermann, Irma Schabussova

**Author notes:** Corresponding authors: Agnieszka Razim and Irma Schabussova. Astrid Laimer-Digruber, Anna M. Schmid, Tanja V. Edelbacher, Magdalena E. Paschal, Tamara Weinmayer, Michael Thaler, Magdalena E. Skalska, Paweł Migdał, Mattia I. Morandi, Catherine Daniel, Dagmar Srutkova, Martin Schwarzer, Jiri Hrdy, Monika Ehling-Schulz, Sabina Górska, Aleksandra Inic-Kanada, Ursula Wiedermann.

## Abstract

Extracellular vesicles (EVs) are key mediators of bacterial communication and adaptation to environmental stress. Their size, cargo, and surface charge are influenced by several factors, including environmental conditions, bacterial physiology, and isolation methods, and are highly strain-specific. Herein, we investigated how exposure to bile, a physiological component of the gut environment, affects the production and properties of EVs released by the probiotic strain *Lactiplantibacillus plantarum* NCIMB 8826. Through ultracentrifugation followed by size-exclusion chromatography (SEC), we isolated highly purified *L. plantarum* EVs (LpEVs) and characterized them according to the Minimal Information for Studies of Extracellular Vesicles guidelines.

SEC purification significantly reduced the contents of contaminating proteins and peptidoglycans, improving the compositional purity of the isolated EVs. Compared with the parent bacteria, purified LpEVs exhibited distinct surface lipid profiles and zeta potential as well as remarkable stability across varying pH levels, elevated NaCl concentrations, and increasing detergent challenges. Under bile stress, the bacteria released larger LpEVs enriched in bile metabolism–related proteins, suggesting vesicle-mediated adaptation. Fourier-transform infrared spectroscopy further revealed bile-induced molecular alterations in LpEVs that differed from those in the parent bacteria.

These findings highlight bacterial EVs as dynamic environmental communicators that respond to stress and may modulate host–microbe interactions before detectable changes occur in the bacterial cells.

## BACKGROUND

Bacterial extracellular vesicles (EVs) are lipid bilayer–enclosed particles secreted by bacteria, typically spherical in shape and ranging within 20–400 nm in diameter [1]. They carry specific cargo such as proteins, lipids, and nucleic acids, with their composition varying depending on whether they originate from gram-positive or gram-negative bacteria [2]. Microbe-associated molecular patterns, recognized by pattern recognition receptors, are present in both the parent bacteria and their EVs. These include lipopolysaccharide (LPS; ligand for Toll-like receptor 4 [TLR4]), lipoproteins (TLR2 ligands), peptidoglycans (PGNs; ligands for nucleotide-binding oligomerization domain–containing proteins NOD1 and NOD2), and nucleic acids such as DNA and small RNAs (TLR7 and TLR9 ligands) [3].

Because of their small size, EVs can interact more readily with host cells than whole bacteria and are capable of disseminating throughout the body [4]. These characteristics make them potent messengers that can modulate host physiological and immune responses. For example, *Escherichia coli* C25–derived EVs induce a dose-dependent inflammatory response in intestinal epithelial cells (IECs) [3], whereas EVs from the probiotic *E. coli* O83 interact with mouse and human airway cells, exerting immunomodulatory effects [5]. Conversely, *Fusobacterium nucleatum* EVs trigger apoptosis in IECs and Caco-2 cells, leading to disruption of epithelial barrier integrity and increased oxidative stress [6]. Additionally, *Vibrio cholerae* EVs have been shown to alter microRNA expression in eukaryotic cells, thereby suppressing host immune responses [7].

The diverse biological activities and molecular composition of bacterial EVs suggest that they fulfill multiple roles in microbial ecology and host interactions. EVs promote colonization through enhanced adhesion, degrade complex polysaccharides, protect bacteria from antibiotics, and facilitate the transfer of antibiotic resistance genes [8]. However, the extent to which EV production is condition-specific remains unclear. In marine bacteria, both the size and rate of EV release can vary within the same strain depending on environmental factors such as nutrient availability, temperature, or light exposure [9]. Under nutrient-deficient conditions, such as those in the undernourished gut, *Bacteroides thetaiotaomicron* EVs scavenge vitamin B12 via high-affinity binding proteins, and these vesicles are selectively internalized by the parent bacteria or host cells [10]. EVs from the same strain can also sequester iron using siderophores produced by other bacteria, thereby reducing iron availability to the host [11].

Despite substantial progress in the study of bacterial EVs, significant knowledge gaps remain. The precise mechanisms by which environmental stressors trigger EV biogenesis and release are not yet fully understood. The evolutionary significance of EV production under stress also remains unclear, particularly regarding how it may confer adaptive advantages in different ecological niches. Therefore, there is an urgent need to develop and refine advanced methodologies for the isolation, characterization, and functional analysis of bacterial EVs not only from artificial culture systems but also under conditions that more closely mimic their natural environments.

In this study, we used *Lactiplantibacillus plantarum* (Lp) NCIMB 8826, a parental strain of Lp WCFS1 originally isolated from human saliva and known for its exceptional adaptive capabilities [12], as a model to investigate the impact of stress on EV production. We isolated Lp EVs (LpEVs) and characterized them according to the Minimal Information for Studies of Extracellular Vesicles (MISEV) guidelines [13]. Size-exclusion chromatography (SEC) was performed for purification, which markedly reduced the contents of contaminating proteins and PGNs. Compared with the parent bacteria, LpEVs exhibited distinct surface lipid profiles and zeta potential (ZP). Stability assays demonstrated that LpEVs were highly resistant to variations in pH, elevated NaCl concentrations, and sodium dodecyl sulfate (SDS) exposure. Notably, LpEVs produced in medium supplemented with 0.25% bile showed increased particle size and protein content, including enrichment in bile metabolism–associated proteins such as bile salt hydrolase (BSH). Fourier-transform infrared (FTIR) spectroscopy further revealed bile-induced spectral changes in LpEVs, reflecting alterations in protein and fatty acid composition under stress. Proteomic and FTIR spectroscopy collectively revealed that exposure to bile induced more pronounced compositional changes in LpEVs than in the parent bacteria. These findings suggest that LpEVs function as indicators of environmental stress, reflecting adaptive responses even before they become detectable at the cellular level.

## METHODS

### EV production and purification

EVs were isolated (**Figure 1**) from Lp [14], kindly provided by Dr. Daniel (Institut Pasteur de Lille, France). Cultures were grown in 5 L of De Man–Rogosa–Sharpe (MRS; Merck, Germany) broth supplemented with Tween 80 (MRS-T) in closed bottles without shaking at 37°C (Innova 4230 Incubator, Eppendorf New Brunswick, Germany). The culture was inoculated with an overnight starter culture at a 1:200 ratio and incubated until an optical density at 600 nm (OD600) of 1.5 was reached. Bacterial cells were then removed via centrifugation (6,000 × *g*, 20 min, 4°C; Heraeus, Thermo Fisher Scientific, USA), followed by filtration through a 0.22-µm sterile vacuum bottle-top filter (Steritop, Merck). The resulting cell-free supernatant was concentrated approximately 13-fold using an Amicon Ultra Stirring Cell (Merck) equipped with 300 kDa ultrafiltration discs.

**Figure 1.**
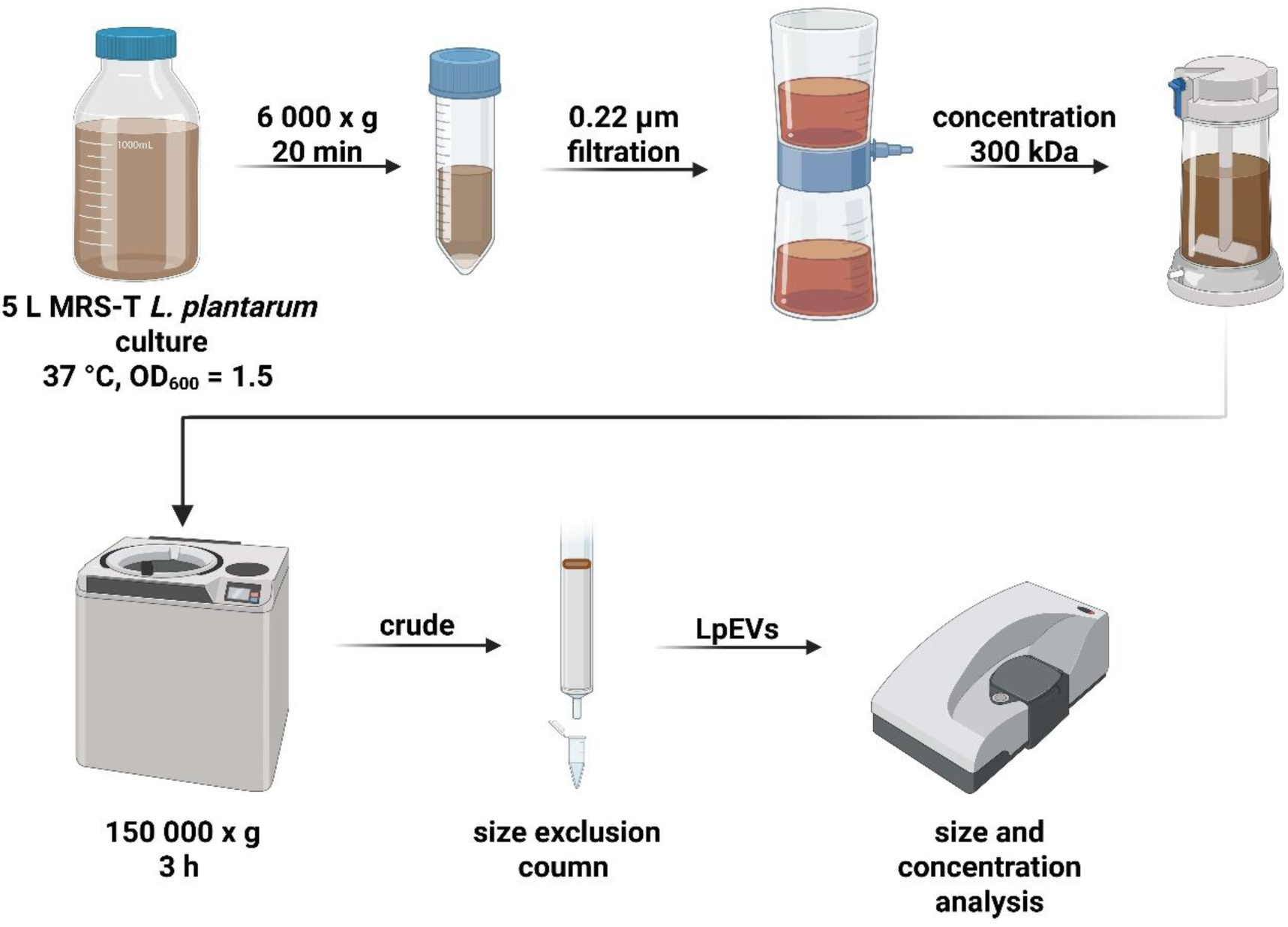
Production and purification of LpEVs. Lp was grown in MRS-T medium at 37°C in a closed bottle until an OD600 of 1.5 was reached. The culture was centrifuged, filtered, concentrated, and subjected to UC. Crude LpEVs were further purified via SEC, pooled, and characterized. Figure created with BioRender.

Crude LpEVs were obtained via ultracentrifugation (UC) of the concentrated supernatant at 150,000 × *g* for 3 h at 4°C (Beckman Coulter, USA; 45Ti rotor) and resuspended in 500 µL of 25 mM 4-(2-hydroxyethyl)-1-piperazineethanesulfonic acid buffer containing 0.15 M NaCl (HEPES-N). The EVs were further purified via SEC using qEVoriginal/35 nm Gen 2 columns coupled to an Automatic Fraction Collector (IZON Science, France), following the manufacturer’s instructions. Eight fractions of 400 µL each were collected (**Figure S1A-B**). Fractions enriched in homogeneous particles—typically corresponding to the purified collection volume (PCV) 1.1–1.3—were pooled and stored at 4°C for immediate use or at −20°C for long-term storage.

MOCK EVs were prepared following the same procedure, using uninoculated 5 L of MRS-T medium. After incubation at 37°C for 8 h (the time required for bacterial cultures to reach an OD600 of 1.5), the medium was processed in parallel with the inoculated samples, including centrifugation, filtration, concentration, UC, SEC, and fraction pooling (PCV 1.1–1.3).

### Characterization of EVs

#### Size, concentration, and ZP measurements

Particle size and concentration were measured using Zetasizer Ultra Red Label (Malvern Panalytical, UK) as previously described [5]. The working ranges were 1 × 10^8^–1 × 1^12^ particles/mL for concentration and 0.3 nm–15 µm for diameter. Analyses were performed in a quartz batch cuvette (ZEN2112, Malvern Panalytical) using General Mode, with all measurements conducted in triplicate. ZP was determined using a disposable folded capillary cell (DTS1070, Malvern Panalytical) [5,15]. EV samples were diluted 1:500 in 1 mM HEPES, whereas bacterial samples from overnight cultures were diluted 1:1000 in 1 mM HEPES. All measurements were performed in triplicate. Data quality and analysis were assessed using Malvern ZS Xplorer software (version 3.2.0.84). Cryogenic electron microscopy (cryo-EM) was additionally employed for EV size validation.

#### Transmission electron microscopy (TEM)

LpEVs were negatively stained as previously described [16]. Briefly, 5 µL of the EV suspension was adsorbed onto formvar- and carbon-coated grids for 20 s, contrasted with 5–10 µL of 1% aqueous uranyl acetate, and blotted twice using the single-drop method. The grids were air-dried and imaged using an FEI Tecnai 20 transmission electron microscope (FEI, Netherlands) equipped with a 4K Eagle CCD camera. Images were processed using Adobe Photoshop.

#### Cryo-EM

For cryo-EM analysis, LpEVs were applied to Quantifoil EM grids (Electron Microscopy Sciences, USA) pre-coated with 10-nm protein A–conjugated colloidal gold particles (Au–NP; Aurion, Netherlands), following an established protocol [5].

### Analysis of surface composition

#### Surface lipids

Surface lipid analysis was performed using time-of-flight secondary ion mass spectrometry (ToF-SIMS) as previously described [5]. Briefly, surface lipids of LpEVs and Lp were analyzed with a ToF-SIMS V instrument (ION-TOF GmbH, Münster, Germany) equipped with a 30 keV Bi3+ ion beam (∼0.61 pA). Positive secondary ions (*m/z* 1–900) were collected from 150 µm × 150 µm areas, and spectra were internally calibrated using standard hydrocarbon peaks. The Bi3+ beam was rastered over a 128 × 128 pixel region, and ion doses were maintained below the static limit (1 × 10^12^ ions/cm^2^) for all analyses. Surface charge neutralization was achieved using a low-energy electron flood gun. The data supporting this analysis are available at RODBUK, Jagiellonian University in Kraków: https://uj.rodbuk.pl/dataset.xhtml?persistentId=doi:10.57903/UJ/GLBXEG.

#### Lipoteichoic acid (LTA)

Lp (10 µg of cell lysate), LpEVs (1.5 × 10^10^ particles), and commercial LTA (10 µg; Sigma-Aldrich, USA) were mixed with 4× Laemmli buffer, boiled for 5 min at 96°C, and separated on a 4%–12% gradient SDS–PAGE gel (Thermo Fisher Scientific, USA). Proteins were transferred onto a polyvinylidene fluoride membrane (Invitrogen, USA) at 30 V for 1 h, blocked with 3% bovine serum albumin (BSA) for 1 h at room temperature (RT), and probed with an anti-LTA antibody (ABAAb02033-1.1, Szabo-Scandic, Austria; 1:500, 2 h, RT). After washing, the membrane was incubated with an alkaline phosphatase–conjugated anti-mouse IgG1 antibody (Sigma-Aldrich, USA; 1 h, RT). The signal was developed using NBT/BCIP (Sigma-Aldrich, USA) and stopped by washing with water. The membrane was imaged using a Gel Doc system (Bio-Rad, USA).

#### PGN

PGN quantification was performed as described by Bito *et al.* [17]. Crude and purified LpEV samples (2.5 × 10^10^ particles in 25 µL) were treated with 100 µL of 1 M NaOH for 30 min at 38°C, followed by acid hydrolysis with 125 µL of 0.5 M H2SO4 and 1,250 µL of concentrated H2SO4. Samples were then boiled for 5 min and rapidly cooled under running water. Subsequently, 12.5 µL of CuSO4 and 25 µL of 1.5% 4-phenylphenol (Sigma-Aldrich, USA) in 96% ethanol were added and incubated for 30 min at 30°C. Absorbance was measured at 560 nm in 96-well plates using a plate reader (Tecan Trading AG, Switzerland).

### Analysis of EV storage stability

The long-term storage stability of LpEVs was evaluated following a modified protocol of Gürgens *et al.* [18]. After UC (150,000 × *g*, 3 h; SW41Ti rotor), LpEVs were resuspended in one of five buffers: (i) 25 mM HEPES with 0.9% NaCl (HEPES-N), (ii) 25 mM HEPES with 0.9% NaCl and 0.2% BSA (HEPES-NA), (iii) phosphate-buffered saline (PBS), (iv) PBS with 0.2% BSA (PBS-A), or (v) PBS with 25 mM HEPES and 0.2% BSA (PBS-HA). Aliquots were prepared in triplicate and stored at 4°C, −20°C, or −80°C. Samples were analyzed at baseline (time 0) and after 1, 6, and 12 months. Each aliquot was thawed only once for size and concentration measurements using the Zetasizer, after which it was discarded. All measurements were performed from a single stock preparation to ensure consistency across analyses.

### Analysis of environmental stability of bacteria and their EVs

The stability of Lp and LpEVs was evaluated under varying salt, pH, and detergent conditions relevant to the human physiological environment.

**Salt stability:** LpEVs (1.2 × 10^11^ particles/mL in 50 µL of HEPES-N) were used as the starting material. Concentrated NaCl was added to achieve final concentrations of 0.25, 0.35, 0.5, and 1.0 M. Samples were mixed and analyzed for particle size and concentration within 30 min. In parallel, overnight bacterial cultures were diluted 1:200 in MRS-T supplemented with the same NaCl concentrations.

**pH stability:** LpEVs (0.5 × 10^10^ particles) were pelleted via UC (150,000 × *g*, 3 h; SW41 rotor), resuspended in 50 µL of HEPES-N adjusted to pH values of 5.7–8.2 (in 0.5-unit increments), and analyzed within 30 min as described above. In parallel, overnight bacterial cultures were diluted 1:200 in MRS-T adjusted to the same pH using citrate–phosphate buffer.

**Detergent stability**: LpEVs (1 × 10^11^ particles/mL in 90 µL of HEPES-N) were incubated with SDS at final concentrations of 0, 0.1, 0.6, 2.2, and 8.9 mM and analyzed within 30 min. In parallel, overnight bacterial cultures were diluted 1:200 in MRS-T containing the same SDS concentrations.

All experiments were repeated three times, and each sample was prepared in triplicate. Bacterial growth under all conditions was monitored by measuring OD600 at 0, 3, 5, 6, 7, and 8 h in 96-well plates using a plate reader. The OD₆₀₀ value at time 0 was used as the baseline and subtracted from subsequent measurements.

### Bile stress assays

#### Bacterial growth and EV production

To evaluate the effect of bile stress, overnight Lp cultures were diluted 1:200 in MRS-T supplemented with 0%–2% (twofold serial dilutions) bovine bile (Sigma-Aldrich, USA). For EV isolation, Lp cultures were grown in 0.25% bovine bile (LpEVs^BILE^), following the same procedure as for standard LpEV preparation. Corresponding MOCK EVs (MOCK EVs^BILE^) were prepared from uninoculated medium processed in parallel.

#### Nucleic acid isolation and analysis

RNA was isolated using an RNA extraction kit (IZON Science, France), and RNase treatment was performed following a previously established protocol [5]. Briefly, 375 µL of LpEVs and LpEVs^BILE^ (∼1.5 × 10^11^ EVs) were treated with proteinase K (100 µg/mL, 2 h, 37°C; Thermo Fisher Scientific, USA), followed by enzyme inactivation at 75°C for 1 h. Subsequently, RNase One (10 U/mL; Promega, USA) was applied for 30 min at 37°C and inactivated with RNase inhibitor (SUPRAase•In^TM^, 1 U/mL; Thermo Fisher Scientific, USA). RNA was extracted in a final volume of 50 µL and analyzed using a Bioanalyzer (Agilent Technologies, USA).

DNA was isolated from 375 µL of the IZON fraction of LpEVs and LpEVs^BILE^ (∼1.5 × 10^11^ EVs) using the FavorPrep Tissue Genomic DNA Extraction Kit (Favorgen Biotech Corporation, Taiwan). DNA concentration was measured using a NanoDrop 1000 spectrophotometer (Thermo Fisher Scientific, USA), and samples were visualized on a 2% agarose gel stained with Midori Green (Thermo Fisher Scientific, USA).

#### Protein quantification and SDS–PAGE

Protein concentrations in LpEVs, LpEVs^BILE^, and the corresponding MOCK controls were determined using three independent assays: (i) Bradford assay (Bio-Rad, USA), (ii) bicinchoninic acid (BCA) assay (Thermo Fisher Scientific, USA), and (iii) Pierce 660 nm Protein Assay (Thermo Fisher Scientific, USA), following the manufacturers’ protocols. Sample volumes were 5 µL (Bradford), 25 µL (BCA), and 10 µL (Pierce). Absorbance was measured using a plate reader. Protein content was calculated as µg of protein per 1 × 10^11^ EVs. BSA standards were prepared in 25 mM HEPES-N. Data obtained from each method were analyzed using one-way analysis of variance (ANOVA), comparing LpEVs^BILE^ with LpEVs. Protein profiles of IZON fractions and pooled samples were assessed via SDS–PAGE using NuPAGE 4%–12% precast gels (Thermo Fisher Scientific, USA), stained with SimplyBlue™ SafeStain (Thermo Fisher Scientific, USA), and visualized using a Gel Doc imaging system (Bio-Rad, USA). A 10–180 kDa PageRuler protein ladder (Thermo Fisher Scientific, USA) was used as a molecular weight reference. A 20 µL aliquot of each sample, corresponding to 1.5 × 10^10^ LpEVs and 0.7 × 10^10^ LpEVs^BILE^, was loaded per lane.

#### Proteomics

Proteins were extracted in SDT buffer (4% SDS, 0.1 M DTT, 0.1 M Tris-HCl, pH 7.6) at 95°C in a thermomixer (750 rpm; ThermoMixer C, Eppendorf, Germany) for 30 min (EVs) or 60 min (bacteria). After centrifugation (20,000 × *g*, 15 min, RT), 5 µg of total protein from each sample was processed using the filter-aided sample preparation method [19] with sequencing-grade trypsin (0.5 µg; Sigma-Aldrich, USA). Resulting peptides were extracted with 2.5% formic acid (FA) in 50% acetonitrile (ACN), followed by extraction with 100% ACN, both solutions containing 0.001% polyethylene glycol [20]. Peptides were then concentrated using a SpeedVac concentrator (Thermo Fisher Scientific, USA).

Liquid chromatography–tandem mass spectrometry was performed using an UltiMate 3000 RSLCnano system (Thermo Fisher Scientific, USA) coupled to a timsTOF Pro mass spectrometer (Bruker Optics GmbH, Germany). Samples were desalted on a trap column (Acclaim PepMap 100 C18, 300-µm ID, 5-mm length, 5-µm particle size; Thermo Fisher Scientific, USA) and washed with 0.1% trifluoroacetic acid. Peptides were eluted in backflush mode from the trap column onto an analytical column (Aurora C18, 75-µm ID, 250-mm length, 1.7-µm particle size; Ion Opticks, Australia) using a 60-min gradient (3%–42% mobile phase B; mobile phase A: 0.1% FA in water; mobile phase B: 0.1% FA in 80% ACN) at a flow rate of 150 nL/min, followed by a system wash with 80% mobile phase B. The trapping and analytical columns were equilibrated before sample injection into the sample loop. The analytical column was installed in the CaptiveSpray ion source (Bruker Optics GmbH, Germany) maintained at 50°C, following the manufacturer’s instructions. The spray voltage and sheath gas were set to 1.5 kV and 1.0 L/min, respectively.

MSn data were acquired in data-independent acquisition (DIA) mode with a base *m/z* range of 100–1700 and a 1/*K0* range of 0.6–1.4 V·s·cm^−2^. The enclosed DIAparameters.txt file defines an *m/z* 400–1000 precursor range with equal window sizes of 26 Th, using two steps for each PASEF scan and a cycle time of 100 ms locked to a 100% duty cycle. High-sensitivity detection mode was applied to measure the EV samples to compensate for the low protein digest input. DIA data were processed in DIA-NN (version 1.8.1) [21] in library-free mode against the cRAP database (http://www.thegpm.org/crap; 111 sequences, version 2018/11) and the UniProtKB protein database for Lp NCIMB 8826 (https://www.uniprot.org/proteomes/UP000000432;

version 2024/01; 3,087 protein sequences). Carbamidomethylation was set as a fixed modification, and trypsin/P was selected as the digestion enzyme, allowing one missed cleavage and peptide lengths of 7–30 amino acids. The false discovery rate threshold was set to 1% at both the precursor and protein levels. MS1 and MS2 accuracies, as well as scan window parameters, were optimized based on initial test searches, using the median parameter values across all samples. Match-between-runs was enabled. Protein MaxLFQ intensities reported in the DIA-NN main output file were further processed using the Omics Workflows containerized environment (https://github.com/OmicsWorkflows; version 4.7.7a).

The downstream data-processing workflow (available upon request) included a) removal of low-quality precursors and contaminant protein groups, b) normalization of precursor intensities using the loessF algorithm, c) calculation and log2 transformation of protein group MaxLFQ intensities, and d) differential expression analysis using the LIMMA statistical test. Proteins with an adjusted *p*-value < 0.05 and a fold change > 2 were considered significantly altered.

The mass spectrometry proteomics data have been deposited in the ProteomeXchange Consortium via the PRIDE partner repository [22] under the dataset identifier PXD060959.

Volcano plots were generated using VolcaNoseR (https://huygens.science.uva.nl/VolcaNoseR/) [23], including proteins identified in at least two samples. Statistical analysis of raw proteomic data was performed using quantitative values processed with the LIMMA statistical package in R. A significance threshold of *p* < 0.05 and a fold-change cutoff of ≤ −2 or ≥ 2 were applied. Manhattan distance was used for ranking significant hits.

STRING analysis in multiple-protein mode was performed on significantly upregulated proteins (LpEVs^BILE^ vs. LpEVs) and proteins uniquely present in LpEVs^BILE^ (904 proteins in total). The Lp WCFS1 protein database was used as the background reference.

#### FTIR spectroscopy

Changes in the metabolic fingerprints of Lp, bile-exposed bacteria (Lp^BILE^), and their corresponding EVs (LpEVs and LpEVs^BILE^) were assessed via FTIR spectroscopy. Three independent cultures were prepared, from which bacterial cells and EVs were collected for matched analyses. Bacterial cells were pelleted (3,000 × *g*, 20 min) and washed with 25 mM HEPES-N. In parallel, EVs from the same cultures were purified as described above. Aliquots (10 µL) of each sample were applied to zinc selenide optical plates (Bruker Optics GmbH, Germany) and dried at 40°C for 30 min. Spectra were recorded in transmission mode using an HTS-XT microplate adapter coupled to a Tensor 27 FTIR spectrometer (Bruker Optics GmbH, Germany) under the following conditions: spectral range of 4000–500 cm^−1^, spectral resolution of 6 cm^−1^, and averaging of 32 interferograms with background subtraction for each spectrum.

Spectra were preprocessed by vector normalization and baseline correction [24,25]. Hierarchical cluster analysis (HCA) was performed on the full spectral range and on regions corresponding to fatty acids (3020–2800 cm^−1^), proteins (1720–1500 cm^−1^), and polysaccharides (1200–900 cm^−1^).

### EV-TRACK

Experimental details have been submitted to the EV-TRACK knowledgebase (EV-TRACK ID: EV250022) [26].

### Statistical analysis

Statistical analyses were performed using GraphPad Prism version 9.5.1, unless otherwise specified. Comparisons between groups were conducted using one-sample *t*-tests, ordinary one-way ANOVA, Dunnett’s multiple-comparisons test with a single pooled variance, or two-way ANOVA, as appropriate. The statistical tests used are indicated in the corresponding figure legends. A *p*-value of ≤0.05 was considered statistically significant.

## RESULTS

### Characterization and stability of LpEVs isolated from Lp cultured in MRS-T

Lp was cultured in MRS-T, and LpEVs were harvested from 5 L cultures at an OD600 of 1.5, corresponding to the mid-logarithmic growth phase. Following filtration to remove cells and debris, crude EVs were isolated via UC. Further purification was performed using SEC (**Figure 1**), yielding pure EVs in eight fractions of 400 µL each (**Figure S1A-B** shows a representative chromatogram). Typically, fractions 1–3 (PCV 1.1–1.3) were pooled, resulting in a final volume of approximately 1.2 mL of purified LpEVs per batch.

Dynamic light scattering (DLS) analysis using a Zetasizer revealed a mean particle size of 73.9 nm (standard deviation [SD] = 6.9 nm) and an average particle concentration of 1.64 × 10^12^ particles/mL (SD = 0.91 × 10^12^ particles/mL) across five independent batches (**Table S1**). The average total yield per 5 L culture was 1.97 × 10^12^ LpEVs (SD = 1.09 × 10^12^). Representative particle size and concentration data from one batch are shown in **Figure 2A-B**. Purification via SEC markedly reduced the protein and PGN content in the purified EV preparations compared with crude EVs obtained via UC alone (**Figure S1C-D**). To contextualize LpEV production relative to bacterial biomass, the colony-forming unit (CFU) count of the parent bacteria was determined in the same 5 L culture used for EV isolation. At an OD600 of 1.5, the culture contained 6.7 × 10^12^ CFUs and yielded 1.2 × 10^12^ LpEVs, corresponding to an approximate bacterium-to-EV ratio of 6:1.

**Figure 2.**
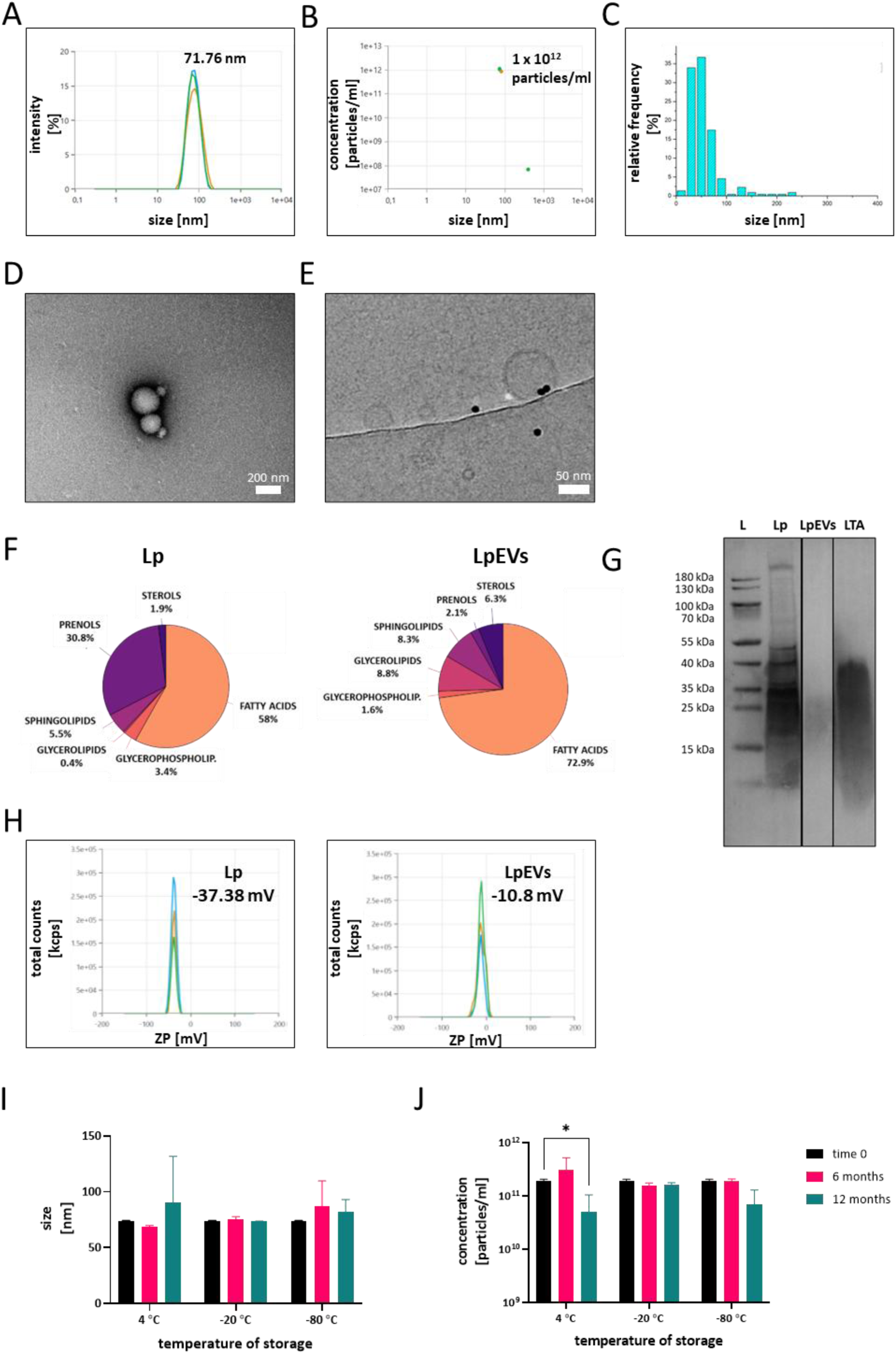
Characterization of LpEVs. Representative measurements of particle size (A) and concentration (B) of LpEVs obtained using a Zetasizer. (C) Size distribution of LpEVs determined using cryo-EM. (D) TEM and (E) cryo-EM visualization of LpEVs. (F) Surface lipid composition of intact Lp and LpEVs analyzed via ToF-SIMS. (G) Western blot analysis of LTA in Lp lysates and LpEVs (L, protein ladder; 10 µg of Lp lysate, 1.5 × 10^10^ LpEVs, and 10 µg of commercial LTA were loaded per lane). (H) ZP measurements of Lp and LpEVs in 1 mM HEPES buffer. Long-term stability of LpEVs stored in 25 mM HEPES-N buffer at 4°C, –20°C, and –80°C showing changes in particle size (I) and concentration (J) over time. Data were analyzed using two-way ANOVA followed by Dunnett’s multiple-comparisons test. Significant differences are indicated as \**p* ≤ 0.05.

Because the determination of EV size largely depends on the analytical method used [5], cryo-EM was employed as a complementary technique to assess the size distribution of LpEVs (**Figure 2C**). Compared with DLS measurements obtained using the Zetasizer, cryo-EM analysis showed that LpEVs predominantly ranged within 20–50 nm in diameter.

TEM (**Figure 2D and S2**) and cryo-EM (**Figure 2E**) were used to visualize the morphology of LpEVs and assess the presence of aggregates or bacterial debris. TEM images showed intact, spherical vesicles that frequently appeared in clusters (**Figure 2D**), as further confirmed by wide-field TEM views (**Figure S2**). Cryo-EM provided definitive confirmation of vesicle identity and revealed a well-defined bilayer membrane even in the smallest EVs (**Figure 2E**), underscoring their structural integrity.

Because MRS-T contains components derived from animal products, it was important to assess whether EVs or EV-like particles originating from the medium itself could confound the experimental results. To address this, MOCK-EVs were isolated from 5 L of uninoculated MRS-T and processed using the same protocol as for bacterial cultures. DLS analysis showed that MOCK-EVs were significantly larger than LpEVs (**Figure S3A**), with a concentration of 2.12 × 10^10^ particles/mL—approximately 100-fold lower than that observed in Lp cultures (**Figure S3B**). TEM revealed only a few particles in MOCK-EV samples, exhibiting morphological features distinct from those of LpEVs (**Figure S3C**). No protein bands were detected when the maximum possible sample volume (20 µL) was loaded onto the SDS–PAGE gel (**Figure S3D**). Moreover, the protein content of the MOCK-EV samples, determined via both BCA and Bradford assays, was below the detection limit (data not shown). These findings confirm that EV-like particles present in MRS-T are negligible in both abundance and protein content and are therefore unlikely to interfere with the interpretation of LpEV-specific experimental results.

The surface lipid composition of Lp and LpEVs was analyzed using the semi-quantitative ToF-SIMS method. The analysis revealed a decrease in prenols and an enrichment of fatty acids on the surface of LpEVs compared with the parent bacteria (**Figure 2F**). Conversely, levels of glycerophospholipids, sphingolipids, and sterols showed only minor differences between the bacteria and their EVs. The presence of LTA, a key structural component of gram-positive bacteria, was confirmed in both the parent cells and LpEVs via western blotting (**Figure 2G**). ZP measurements performed in 1 mM HEPES buffer revealed distinct differences in surface charge between the bacteria and their EVs (**Figure 2H**). The bacterial surface exhibited a strong negative charge of –37.38 mV (SD = 0.25 mV), whereas LpEVs displayed a considerably less negative charge of –10.8 mV (SD = 1.01 mV). These results indicate substantial differences in the surface properties of LpEVs and their parental cells, which may critically influence their stability, interactions, and biological functionality.

LpEVs were routinely stored in HEPES-N buffer at –20°C. To evaluate their long-term stability under different conditions, we analyzed LpEV samples stored in HEPES-N at three temperatures: 4°C, –20°C, and –80°C (**Figure 2I-J**). At 4°C, signs of instability became evident after 12 months, including significant fluctuations in particle concentration. Conversely, samples stored at –20°C and –80°C remained stable for up to one year. We further tested four alternative buffer formulations for their ability to preserve LpEVs: (i) PBS, (ii) PBS-A, (iii) PBS-HA, and (iv) HEPES-NA (**Figure S4**). At 4°C, none of the buffers maintained LpEV stability beyond six months; complete degradation occurred, as no reliable measurements could be obtained (low data quality reported by the Zetasizer) (**Figure S4A**). Conversely, storage at – 20°C (**Figure S4B**) and –80°C (**Figure S4C**) preserved vesicle integrity across all buffer conditions. Notably, buffers containing albumin led to an apparent increase in particle size, suggesting aggregation or interaction effects at 4°C. Based on these results, we recommend storing LpEVs at –20°C in 25 mM HEPES-N buffer as the optimal condition for maintaining vesicle integrity and concentration during long-term storage.

### LpEVs retain stability under variable pH, high salt, and low SDS concentrations

Given the apparent robustness of LpEVs, we evaluated their stability under various environmental conditions that mimic those in the human body. Equivalent quantities of LpEVs were exposed to buffers of different compositions, and their particle size and concentration were measured using the Zetasizer. LpEVs remained stable across a pH range of 5.7–7.7, showing no significant changes in particle size or concentration. However, at pH values above 7.7, a significant decrease in LpEV concentration was observed, indicating sensitivity to alkaline conditions (**Figure 3A**). LpEVs also demonstrated high tolerance to salt stress: incubation in NaCl concentrations up to 1 M caused no notable alterations in particle size or concentration (**Figure 3B**). When exposed to the negatively charged detergent SDS, LpEVs maintained stability at concentrations up to 0.6 mM. At higher SDS concentrations, both particle size and concentration changed significantly; however, LpEVs remained detectable even under these conditions, suggesting that complete vesicle degradation did not occur (**Figure 3C**).

**Figure 3.**
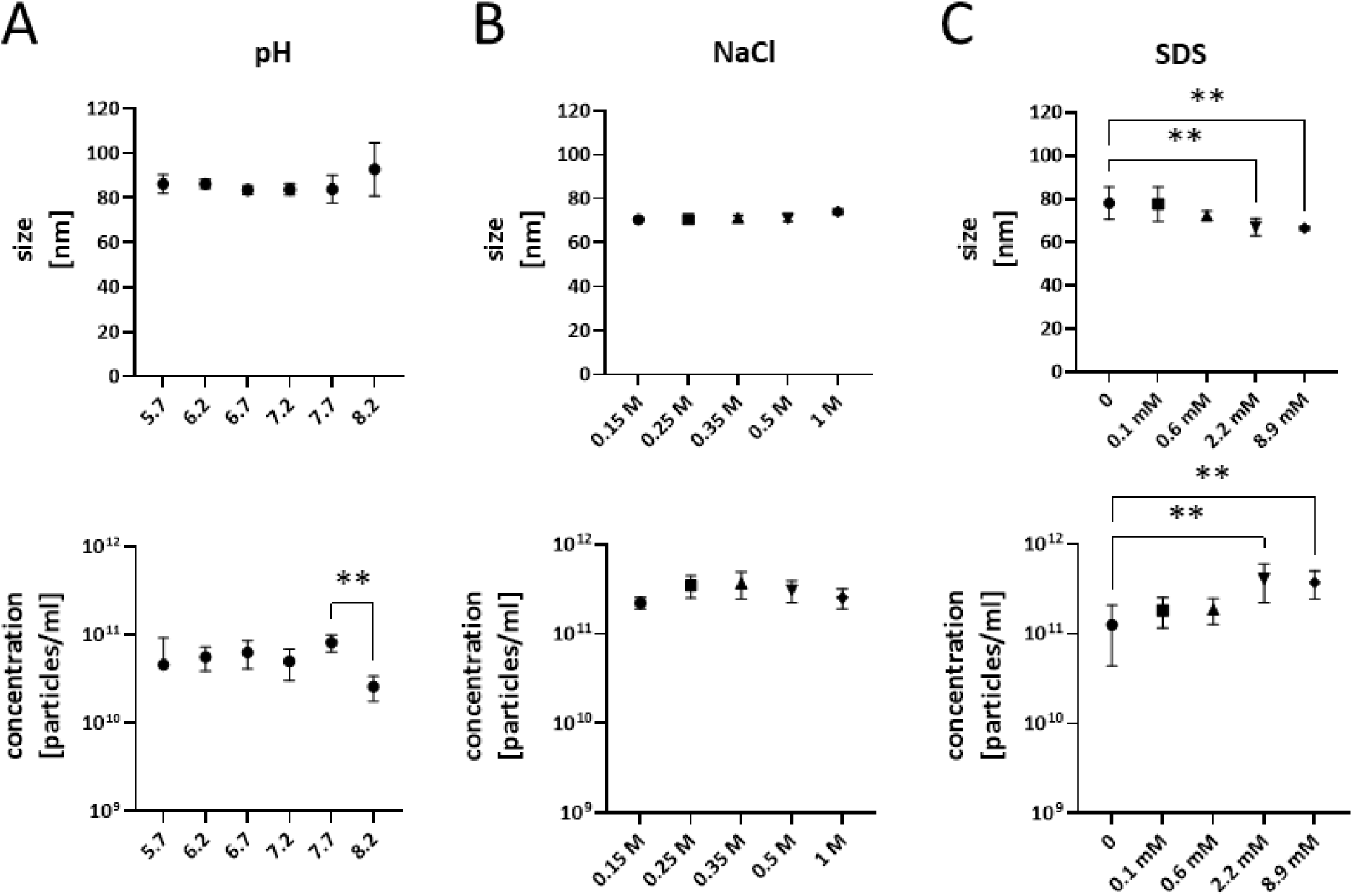
Influence of pH, salt, and detergent on the stability of LpEVs. Particle size and concentration of LpEVs were measured after exposure to different pH values (A), NaCl concentrations (B), and SDS concentrations (C) using a Zetasizer. Data were analyzed using one-way ANOVA followed by Dunnett’s multiple-comparisons test. Significant differences are indicated as \*\**p* ≤ 0.01.

### Growth of Lp is significantly impaired by environmental stressors

Given the demonstrated stability of LpEVs across a broad range of pH values, salt concentrations, and negatively charged detergents, we next examined how these environmental conditions affect the growth of the parent bacteria. Lp was initially cultured in MRS-T, and overnight cultures were subsequently inoculated into fresh media modified to reflect varying pH, NaCl, and SDS concentrations (**Figure 4**). Under standard conditions (pH 5.8), the bacteria exhibited robust growth. However, deviations from this pH significantly impaired bacterial proliferation, with significant reductions observed from 5 h post-inoculation (*p* < 0.0001, two-way ANOVA) (**Figure 4A**). Similarly, increasing the NaCl concentration above 0.15 M led to dose-dependent growth inhibition, with significant differences detected at all tested concentrations from 6 h onward (*p* < 0.0007, two-way ANOVA) (**Figure 4B**). Exposure to SDS also markedly suppressed bacterial growth; even at 0.6 mM, SDS significantly reduced growth at 5, 6, 7, and 8 h post-inoculation (*p* < 0.0001, two-way ANOVA), with higher concentrations exerting progressively stronger inhibitory effects (**Figure 4C**).

**Figure 4.**
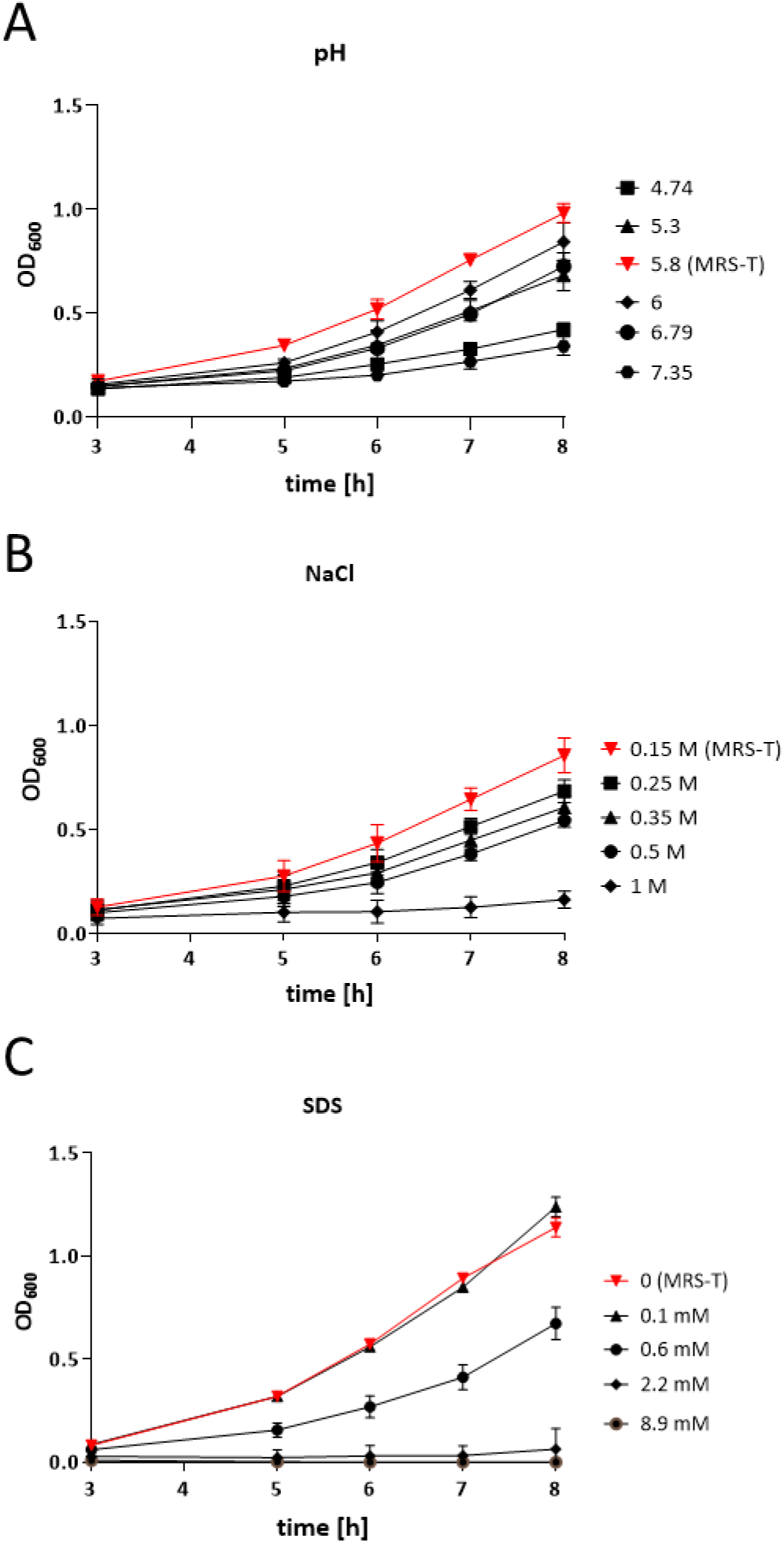
Influence of pH, salt, and detergent on Lp growth. OD600 measurements of Lp cultures grown in media (A) adjusted to pH values ranging within 4.74–7.35 (pH measured at the end of culture), (B) supplemented with NaCl concentrations of 0.15–1.0 M, and (C) supplemented with SDS concentrations of 0–8.9 mM. Data were analyzed using two-way ANOVA followed by Dunnett’s multiple-comparisons test (see statistical analysis in the text).

### Lp produces distinct EV populations in response to bile stress

As Lp encounters bile salts during gastrointestinal transit, we investigated bile as a physiologically relevant stressor influencing bacterial growth, EV production, and EV-associated cargo. As shown above, environmental factors such as pH, salinity, and detergents affect bacterial growth dynamics. Extending this to bile, we examined the effects of increasing concentrations (0%–2%) of bovine bile in MRS-T. A concentration of 0.25% bile was selected for detailed analysis, as it caused approximately a 50% reduction in bacterial growth (**Figure 5A**), indicating substantial physiological stress. EVs produced under bile stress (LpEVs^BILE^) were isolated and characterized using the same protocol as for EVs produced under standard conditions (LpEVs), with harvesting conducted at OD600 = 1.5. Although the total particle yield of LpEVs^BILE^ was comparable to that of LpEVs, DLS analysis revealed a significant increase in mean vesicle size, from 82.79 nm (SD = 11.63 nm) to 118.1 nm (SD = 8.35 nm) (**Figure 5B**). Additionally, ZP measurements showed a more negative surface charge for LpEVs^BILE^ (–17.09 mV, SD = 1.33 mV) than for LpEVs (–10.99 mV, SD = 2 mV), suggesting alterations in membrane composition or structure (**Figure 5C**). Analysis of vesicular RNA content using a Bioanalyzer revealed a significant enrichment of RNA in LpEVs^BILE^, increasing from 2.21 ng per 1 × 10^11^ EVs in LpEVs to 155.56 ng per 1 × 10^11^ EVs in LpEVs^BILE^. Following RNase treatment, both vesicle types retained small RNA fragments ≤100 nucleotides, whereas untreated samples (particularly LpEVs) contained a broader range of RNA sizes, including longer fragments (**Figure 5D**). No DNA was detected in either EV preparation (data not shown).

**Figure 5.**
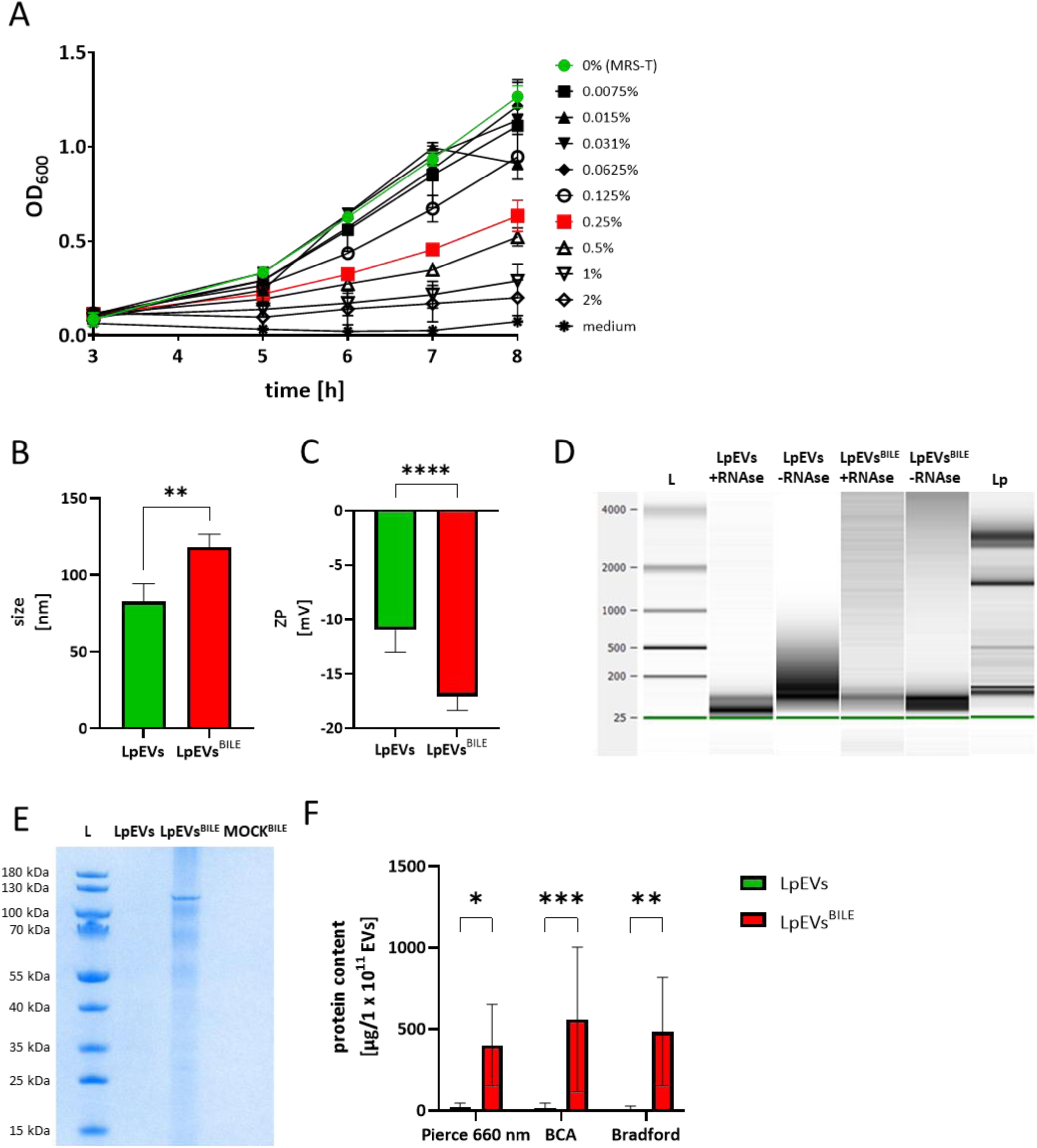
Influence of bile on the growth of Lp and the characteristics of its EVs. (A) Growth curves of Lp cultured in MRS-T supplemented with increasing concentrations of bovine bile (0%–2%). (B) Size distribution and (C) ZP measurements of LpEVs and LpEVs^BILE^ obtained using a Zetasizer. (D) RNA content in untreated and RNase-treated EVs analyzed using a Bioanalyzer. L, RNA ladder. (E) SDS–PAGE of equal EV inputs (based on particle number) from LpEVs and LpEVs^BILE^, and 20 µL of MOCK-EVs^BILE^. L, protein ladder. (F) Total protein content of LpEVs and LpEVs^BILE^ quantified using the Pierce 660 nm Protein Assay, BCA, and Bradford assays. Results are normalized to µg of protein per 1 × 10^11^ EVs and expressed as mean ± SD. Panels B and C were analyzed using unpaired two-tailed *t*-tests; panel F was analyzed using two-way ANOVA. Significance levels are indicated as *p* < 0.05, \**p* < 0.01, \*\**p* < 0.001, and \*\*\**p* < 0.0001.

To investigate the protein cargo of LpEVs and LpEVs^BILE^, we performed SDS–PAGE, quantitative protein assays using three independent methods, and subsequent proteomic analysis. SDS–PAGE revealed a distinct protein band at approximately 120 kDa in LpEVs^BILE^, whereas no visible bands were detected in LpEVs under identical loading conditions or in the MOCK-EVs^BILE^ sample (**Figure 5E**). Given the typically low protein yield from bacterial EVs—particularly for LpEVs—this likely reflects protein concentrations below the detection limit of SDS–PAGE rather than a true absence of protein. Consistently, quantitative protein assays showed a significantly higher total protein content in LpEVs^BILE^ than in LpEVs (**Figure 5F**), with no significant variation among the three methods used.

### FTIR spectroscopy revealed bile-induced spectral alterations in Lp and its EVs

FTIR spectroscopy was employed to investigate bile-induced metabolic changes in Lp (**Figure 6A**) and its EVs (**Figure 6B**), building on previous studies demonstrating the utility of this technique for analyzing biochemical profiles of cells and vesicles [24,25]. Spectral data were recorded in the 4000–500 cm^−1^ range and preprocessed before analysis. The effect of bile on LpEV spectra was evident across all three major spectral regions—proteins (1720–1500 cm^−1^), fatty acids (3020–2800 cm^−1^), and polysaccharides (1200–900 cm^−1^)—but was most pronounced in the protein region (**Figure 6B**). A distinct new peak appeared at 1550 cm^−1^ in LpEVs^BILE^ that was absent in LpEVs (indicated by an arrow in **Figure 6B**). This wavelength is characteristic of chemical groups containing amide bonds, carboxylates, or nitro compounds. In contrast, spectral differences in the bacterial samples were primarily observed in the regions associated with fatty acids and polysaccharides (**Figure 6A**). Baseline- and vector-normalized spectra from two additional biological replicates of bacteria and EVs are provided in the Supplementary Data (**Figure S5A-B**).

**Figure 6.**
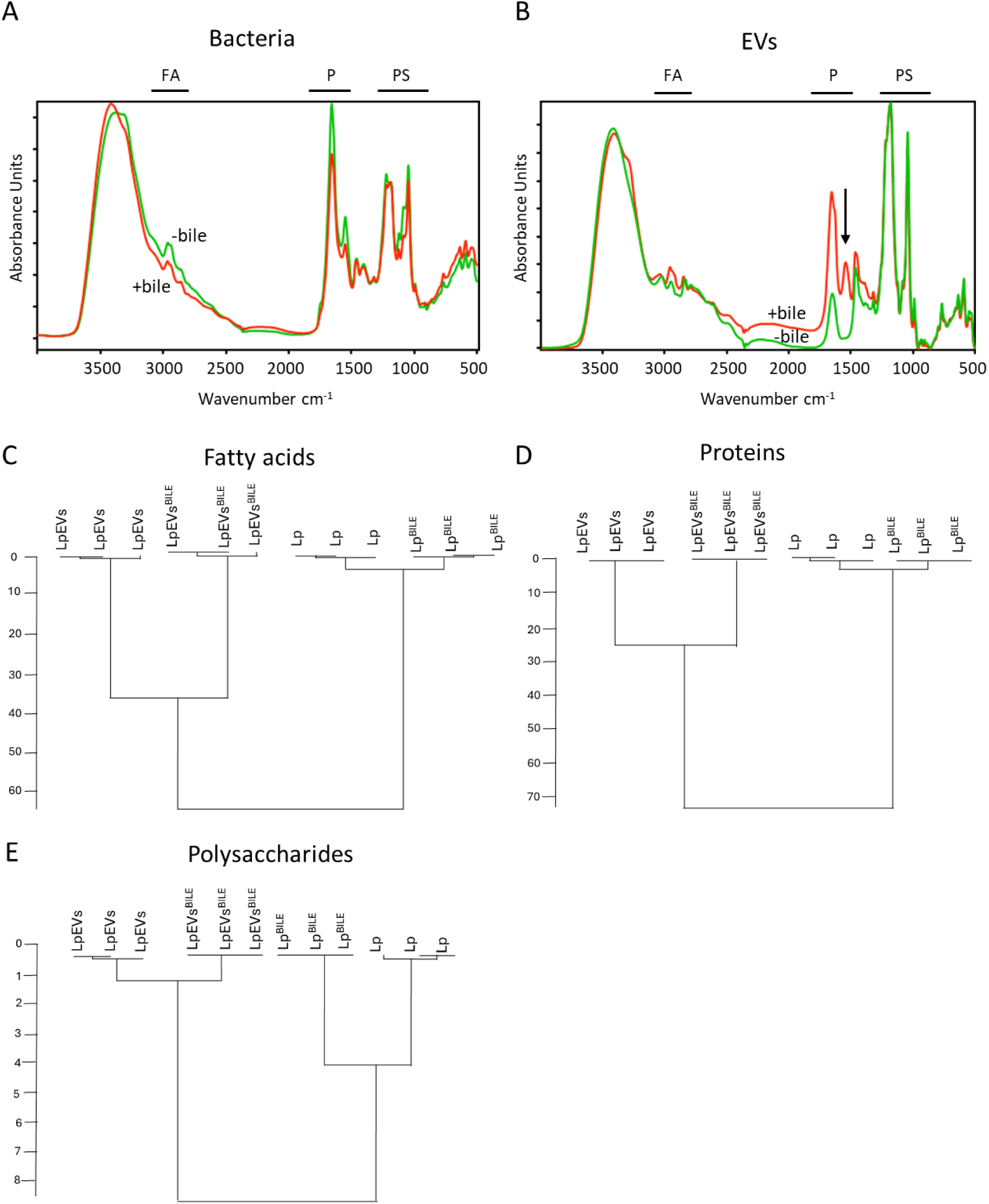
Metabolic fingerprints of Lp, Lp^BILE^, LpEVs, and LpEVs^BILE^ obtained via FTIR spectroscopy. FTIR spectral profiles of Lp, Lp^BILE^, LpEVs, and LpEVs^BILE^ were recorded and analyzed. Representative spectra for the parent bacteria (A) and EVs (B) are shown, with specific regions indicated: FA, fatty acids; P, proteins; PS, polysaccharides. HCA was performed on preprocessed and vector-normalized spectra using the Ward algorithm at repro level 30, based on spectral regions corresponding to fatty acids (C), proteins (D), and polysaccharides (E). Data represent three biological replicates, each measured in triplicate (technical replicates).

To further investigate the impact of bile on the spectral profiles of Lp and LpEVs, HCA was performed. This analysis revealed clear clustering of samples according to the presence or absence of bile in the culture medium, whether based on the entire spectral range (**Figure S5C**) or specific regions: fatty acids (**Figure 6C**), proteins (**Figure 6D**), and polysaccharides (**Figure 6E**). Although the overall impact of bile on the spectral heterogeneity of Lp/Lp^BILE^ and LpEVs/LpEVs^BILE^ was comparable when analyzed across the full spectrum (**Figure S5C**), region-specific differences became apparent (**Figure 6C-E**). Spectral heterogeneity between LpEVs and LpEVs^BILE^ was more pronounced in the fatty acid and protein regions, whereas bacterial cells exhibited greater variability in the polysaccharide region.

### LpEVs^BILE^ show higher detection and relative abundance of bile salt hydrolase and catalytic enzymes

To investigate further qualitative differences in protein composition, we performed a comparative proteomic analysis of Lp cultures grown with and without 0.25% bile, along with the corresponding EVs produced under these conditions. The proteomic workflow demonstrated excellent reproducibility across biological and technical replicates (**Figure S6A**). Consistent with the quantitative protein data (**Figure 5F**), LpEVs^BILE^ contained significantly higher total protein content and a greater number of identifiable proteins than LpEVs (**Figure S6B**). For mass spectrometry, 250 ng of trypsin-digested peptide was analyzed for Lp, Lp^BILE^, and LpEVs^BILE^, whereas the maximum possible volume of LpEVs was used due to their limited protein content. In bacterial cells, a total of 1,871 proteins were identified in both Lp and Lp^BILE^, with only 56 showing altered abundance (**Figure 7A**). Among these, BSH, a key enzyme mediating bile acid deconjugation [27], showed no significant change. Seventy-one proteins were unique to Lp, whereas 22 were detected exclusively in Lp^BILE^. In contrast, the EV proteome exhibited substantially greater variation. Across LpEVs and LpEVs^BILE^, 1,627 proteins were identified, with 101 showing significant differential abundance (**Figure 7B**). Notably, BSH was significantly upregulated in LpEVs^BILE^ compared with LpEVs. Furthermore, 82 proteins were unique to LpEVs, whereas 784 were detected exclusively in LpEVs^BILE^, indicating that bile stress induces a pronounced shift in EV cargo composition.

**Figure 7.**
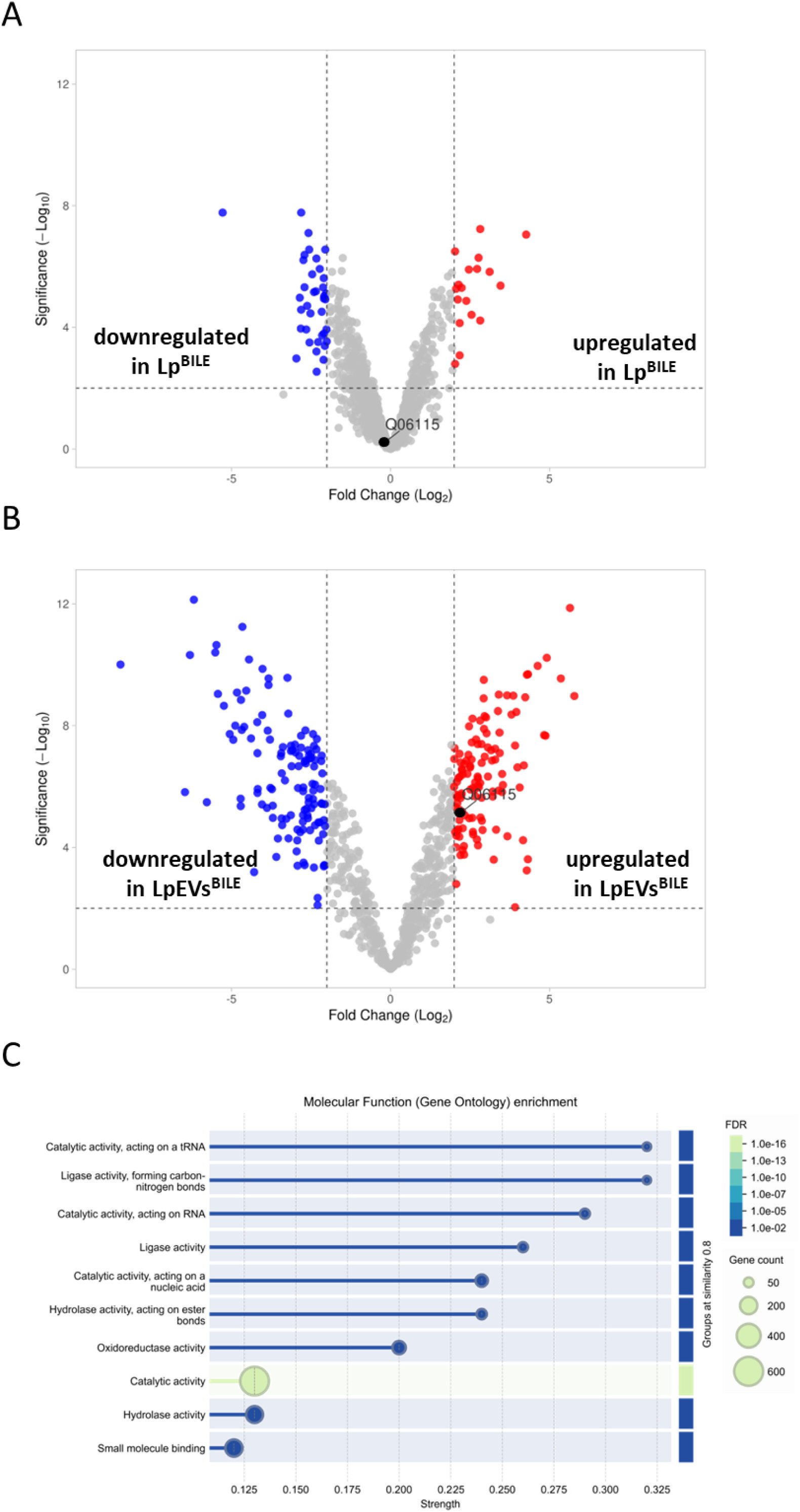
Proteomic analysis of Lp, Lp^BILE^, LpEVs, and LpEVs^BILE^. (A) Volcano plot showing differential protein abundance between Lp cultured without and with bile. BSH (Q06115) is highlighted with a black dot. (B) Volcano plot showing differences in the protein profiles of LpEVs^BILE^ compared with LpEVs. BSH is highlighted with a black dot. (C) Gene Ontology enrichment analysis of molecular functions in LpEVs^BILE^ compared with the Lp reference proteome, performed using STRING.

Protein–protein interaction network and functional enrichment analyses performed using STRING revealed significant enrichment of proteins associated with molecular functions such as catalytic, ligase, hydrolase, and oxidoreductase activities (**Figure 7C**). Biological processes significantly enriched in LpEVs and LpEVs^BILE^ included the metabolism of pyridine-containing compounds, tRNA and RNA processing, noncoding RNA processing, and vitamin metabolic pathways (**Table S2**). Compared with the bacterial proteome background, LpEVs^BILE^ were predominantly enriched in cytosolic and cytoplasmic proteins (**Table S2**). Keyword-based annotation further revealed enrichment in flavoproteins, tRNA processing, ligases, oxidoreductases, transcription-associated proteins, and cytoplasmic proteins (**Table S2**). Of the 30 flavoproteins identified in Lp, 24 were also detected in LpEVs^BILE^.

## DISCUSSION

Probiotics are live microorganisms that confer health benefits to the host when administered in adequate amounts [28]. These beneficial microbes support various physiological functions, both directly and indirectly, including protection against pathogens and attenuation of inflammation [29]. To exert these effects, probiotics must survive transit through the gastrointestinal tract, specifically by withstanding gastric acid and bile salts, and reach the intestine in sufficient numbers [30]. We previously demonstrated that Lp exhibits remarkable resilience under harsh conditions [31]. This resilience enables it to persist within the host environment, exerting both local immunomodulatory effects at mucosal surfaces and systemic benefits [32,33]. More recently, bacterial EVs have been recognized as key mediators in this process, providing a cell-free mechanism through which probiotics enhance host–microbe communication and contribute to environmental resilience [34].

In this study, we provide new insights into the production, purification, characterization, and environmental adaptability of EVs from Lp NCIMB 8826, originally isolated from human saliva. Through differential UC combined with SEC, and following the MISEV guidelines [13], we obtained highly purified LpEVs with consistent physicochemical properties across multiple batches. The ability to produce EVs appears to be a conserved trait among Lp strains, as isolates from fermented tea [35], various foods [36], and kimchi [37,38] have also been reported to release EVs, highlighting the ecological versatility and potential functional relevance of EV production.

Our results also underscore the critical importance of the purification strategy. SEC substantially reduced the contents of co-isolated contaminants such as PGN and nonvesicular proteins, consistent with a previous report [36]. PGN, a key component of bacterial cell walls, is essential for maintaining structural integrity and resisting internal turgor pressure [39]. Although PGN was detected in the crude LpEV preparation, as also observed by Morishita *et al.* [40], it was largely removed via SEC, supporting the interpretation that PGN represents a co-purified contaminant rather than a genuine EV cargo. This finding aligns with our previous observations in *E. coli* O83, where extracellular components were shown to co-purify with EVs and potentially confound functional analyses [5]. Because crude and purified EVs may exert distinct biological effects due to such contaminants, implementing rigorous and standardized EV isolation protocols is essential to enable meaningful comparisons across studies. Moreover, this raises an important question: do highly purified EVs truly reflect their physiological counterparts within the mammalian host? *In vivo*, EVs are likely to acquire a biomolecular corona—a layer of loosely or tightly associated host and environmental molecules—that can modulate their biological activity [41]. Although this phenomenon is well documented for mammalian EVs [42,43], the existence and functional relevance of a corona on bacterial EVs remain largely unexplored. Addressing this knowledge gap will be crucial for balancing methodological stringency with biological relevance, ensuring that functional studies capture the true complexity of EV behavior within the host environment.

Although optimizing isolation techniques is essential for obtaining high-purity EVs, they are not the sole determinants of EV composition. Biological and environmental factors—including temperature [9], growth medium composition [44,45], pH [46], culture duration [47,48], and strain-specific characteristics [49]—can independently influence EV yield, physicochemical properties, and immunomodulatory potential.

The biophysical properties of EVs also have functional significance. ZP analysis revealed that LpEVs possessed a lower surface charge than their parent cells, consistent with previous findings by Rogers *et al.* [15], who reported that EVs often differ in surface charge from their parental bacteria. Such differences reflect variations in membrane composition and may influence vesicle stability, host interactions, and environmental adaptability.

Bacterial EVs also show great potential as vaccine platforms [50]. The clinical success of Bexsero—a licensed vaccine based on *Neisseria meningitidis* EVs—demonstrates the feasibility of translating bacterial EVs into effective immunotherapies [51]. However, to advance EV-based vaccines from bench to bedside, it is crucial to ensure their stability during storage and their resilience under physiologically relevant conditions [18]. We found that LpEVs are remarkably robust across a range of environmental stresses, including variations in pH, salt concentration, detergent exposure, and long-term storage at 4°C, making them promising candidates for scalable therapeutic applications.

Bacterial EVs are increasingly recognized as universal mediators of microbial and interspecies communication [33,34]. Owing to their small size and structural robustness, they can translocate from the gut into the portal circulation and deliver molecular cargo to peripheral sites [35,36]. Among these cargo components, RNA has emerged as a key determinant of functional relevance. In our study, LpEVs derived from both bile-treated and untreated cultures contained diverse RNA populations, including small RNAs (sRNAs) protected from RNase digestion. Supporting their functional potential, Yu *et al.* demonstrated that sRNAs in EVs from the same Lp strain can mediate interspecies signaling, with sRNA71 shown to suppress gene expression in human cells [52]. Interestingly, we observed an increased total RNA yield in LpEVs following bile exposure, suggesting that environmental conditions influence vesicular RNA content, possibly through a selective packaging mechanism. This observation aligns with previous findings in *Staphylococcus aureus*, wherein vesicular RNA profiles varied with growth phase and antibiotic stress [53]. Whether bile-induced sRNAs differ functionally from those in untreated EVs remains to be investigated.

Bile, a natural physiological stressor, regulates intestinal microbial populations by inhibiting excessive bacterial growth, thereby driving gut bacteria to evolve adaptive survival mechanisms [54]. Species such as *Lactobacillus* and *Bifidobacterium* respond to bile exposure by reinforcing their S-layer and increasing the production of polysaccharides, bile-responsive transporters, and enzymes such as BSH, which deconjugates bile acids. Beliakoff *et al.* demonstrated that bile modulates gene transcription in *Lactobacillus johnsonii*, particularly affecting genes involved in transport, biosynthesis, and cell wall processes during the late exponential phase [45]. In our study, which focused on Lp during the logarithmic growth phase and on proteins regulated post-transcriptionally, we observed no substantial changes in the overall bacterial protein content in response to bile exposure. Neither proteomic analysis nor FTIR spectral profiling revealed significant alterations in protein expression or bacterial biochemical fingerprints. These findings are consistent with those of Koskenniemi *et al.*, who reported only minor proteomic changes after 60 min of bile exposure, suggesting that the activation of adaptive mechanisms may require a longer period to manifest [55].

In our study, Lp produced significantly larger EVs in response to bile exposure, without a corresponding increase in vesicle number. This contrasts with previous observations in *L. johnsonii* N6.2, wherein bile treatment increased both EV size and abundance [45]. The discrepancy in vesiculation patterns may reflect species-specific differences in membrane composition, EV biogenesis pathways, or stress-response mechanisms activated by bile. One possible explanation is that Lp exerts stricter control over membrane remodeling under bile stress. Alternatively, bile may promote vesicle enlargement through membrane fusion or altered budding dynamics, independent of vesicle yield. Overall, these findings underscore the influence of strain-specific physiological traits on EV production under environmental stress.

Interestingly, exposure to bile also altered the surface charge of LpEVs, as indicated by a marked reduction in ZP. A more negatively charged surface could influence vesicle stability, colloidal behavior, and interactions with host cells or mucosal surfaces. Similar effects have been reported in *Bifidobacterium* species, wherein the steroid moieties of bile salts interact with hydrophobic domains on the bacterial surface [56]. It is plausible that analogous interactions occur at the LpEV membrane, thereby modulating its surface charge and molecular composition. Supporting this hypothesis, LpEVs^BILE^ consistently exhibited a yellow coloration, contrary to the colorless appearance of control vesicles, suggesting direct incorporation or surface association of bile components (data not shown). To sup up, given the presence of bile salts, and also Tween 80 in the growth medium, matrix contributions to the size and charge changes cannot be excluded.

FTIR spectroscopy provides a powerful approach for detecting metabolic and compositional changes in bacterial EVs in response to environmental stressors. Although FTIR spectroscopy has previously been applied to fingerprint *Bacillus cereus* and its EVs [25], our study extends its utility by demonstrating its sensitivity to bile-induced alterations. Specifically, FTIR spectroscopy revealed significant alterations in spectral regions corresponding to proteins and fatty acids. Of particular interest was the appearance of a peak at 1550 cm^−1^ in LpEVs^BILE^ compared with control EVs, consistent with the characteristic infrared absorbance of the isoalloxazine ring in flavin cofactors [57]. Given the established role of flavoproteins in bile metabolism [58], this spectral feature may indicate the selective incorporation of flavin-dependent enzymes into EVs in response to bile exposure.

Further proteomic analysis revealed significant compositional changes in LpEVs^BILE^: 101 proteins were differentially abundant, whereas 784 were uniquely detected compared with control LpEVs. Notably, enzymes with catalytic, hydrolytic, and ligase activities—including BSH—were enriched, suggesting effects on lipid, polysaccharide, and bile acid metabolism. This functional signature may reflect adaptive responses to bile stress or suggest regulated cargo selection, whereby Lp selectively packages proteins into EVs under specific environmental conditions.

A methodological limitation should be noted: LpEVs produced without bile acid yielded low peptide mass, necessitating injection of the full available sample rather than a fixed protein amount. This increases data missingness and may accentuate apparent enrichment patterns. Nonetheless, our findings are align with those of Stentz *et al.*, who demonstrated that *B. thetaiotaomicron* produces EVs enriched in proteins linked to bile acid metabolism *in vivo* [59]. Such stress-responsive sorting mechanisms may promote bacterial survival or modulate host physiology through EVs.

The FTIR spectra of Lp showed also more subtle changes, primarily in the polysaccharide region following bile exposure, supporting a previously reported finding that stress can alter polysaccharide content [60]. Proteomic analysis further corroborated this stress-adaptive response. Notably, the transcriptional repressor LsrR, which negatively regulates polysaccharide biosynthesis [61], was consistently detected in LpEVs^BILE^ but was absent in control LpEVs. This selective packaging suggests that Lp may employ EVs to export regulatory proteins such as LsrR, potentially modulating gene expression and contributing to stress adaptation. In parallel, bile exposure upregulated several stress-response regulators in both bacterial cells and EVs, including Gls24 family homologs Asp1 and Asp2, and the small heat shock protein Hsp3. These proteins are associated with membrane stabilization, protein folding, and general stress resilience, underscoring the coordinated physiological adaptation of Lp to bile challenge. Overall, this study highlights the value of integrating FTIR spectroscopy–based metabolic profiling with proteomic analysis to elucidate bile-induced biochemical dynamics in both bacterial cells and their EVs. Together, these findings suggest that EVs may serve as vehicles for exporting regulatory proteins, thereby contributing to metabolic reprogramming and stress adaptation.

## CONCLUSIONS

Overall, our findings highlight the remarkable adaptability of gut bacteria such as Lp, which utilize EV production to dynamically respond to environmental challenges. The apparent stimulus-specific nature of EV cargo sorting warrants further investigation. Comparative studies involving other stressors—such as nutrient limitation or oxidative stress—will be essential to determine whether the bile-specific EV responses observed here represent a broader adaptive strategy. Elucidating these mechanisms will be crucial for advancing the therapeutic application of bacterial EVs.

## Supporting information

Information on the access to databases

## LIST OF ABBREVIATIONS

EVs: extracellular vesicles
SEC: size-exclusion chromatography
LpEVs: *L. plantarum* EVs
TLR4: Toll-like receptor 4
PGNs: peptidoglycans
IECs: intestinal epithelial cells
Lp: *Lactiplantibacillus plantarum*
MISEV: Minimal Information for Studies of Extracellular Vesicles
ZP: zeta potential
SDS: sodium dodecyl sulfate
FTIR: Fourier-transform infrared
MRS: De Man–Rogosa–Sharpe
MRS-T: MRS broth supplemented with Tween 80
OD600: optical density at 600 nm
UC: ultracentrifugation
HEPES: 4-(2-hydroxyethyl)-1-piperazineethanesulfonic acid
PCV: purified collection volume
cryo-EM: cryogenic electron microscopy
TEM: transmission electron microscopy
ToF-SIMS: time-of-flight secondary ion mass spectrometry
LTA: lipoteichoic acid
BSA: bovine serum albumin
RT: room temperature
PBS: phosphate-buffered saline
LpEVs^BILE^: bile-exposed LpEVs
mock EVs^BILE^: bile-exposed mock EVs
BCA: bicinchoninic acid
ANOVA: analysis of variance
FA: formic acid
ACN: acetonitrile
DIA: data-independent acquisition
Lp^BILE^: bile-exposed bacteria
HCA: hierarchical cluster analysis
DLS: dynamic light scattering
SD: standard deviation
CFU: colony-forming unit
BSH: bile salt hydrolase
sRNAs: small RNAs

## DECLARATIONS

### Ethics approval and consent to participate

Not applicable

### Consent for publication

Not applicable

### Availability of data and materials

The authors have deposited the following datasets in open repositories: ProteomeXchange Consortium via the PRIDE partner repository (dataset identifier PXD060959) and RODBUK Jagiellonian University in Kraków https://uj.rodbuk.pl/dataset.xhtml?persistentId=doi:10.57903/UJ/GLBXEG;

DOI: 10.57903/UJ/GLBXEG). Additional data are available from the corresponding author upon reasonable request.

### Authors’ contributions

A.R. - investigation, funding acquisition, conceptualization, methodology, visualization, writing - review & editing, writing - original draft, data curation. A.L.D. - investigation, methodology, visualization, writing - original draft, writing - review & editing. A.M.S. – investigation, methodology. T.V.E. – investigation. M.E.P. - investigation, methodology. M.T. – investigation. M.E.S. - investigation, methodology, data curation. P.M. - investigation, methodology. M.M. - investigation, methodology. C.D. - resources, writing - review & editing. D.S. - investigation, writing - review & editing. M.S. - investigation, writing - review & editing. J.H. - investigation, writing - review & editing. M.E.S. - supervision, methodology, writing - review & editing. S.G. - supervision, methodology, writing - review & editing. A.I.K. - methodology, investigation, writing - review & editing. U.W. – resources. I.S. - funding acquisition, conceptualization, writing - original draft, writing - review & editing, supervision, resources, data curation, project administration. All authors read and approved the final manuscript.

## Acknowledgments

The authors acknowledge Enago (www.enago.com) for providing professional English language editing and formatting services in the preparation of this manuscript for submission.

## Authors’ information

AR - trained biotechnologist and Assistant Professor at the Hirszfeld Institute of Immunology and Experimental Therapy, Polish Academy of Sciences, and co-founder of Micronose. AR has extensive research experience in probiotics, host–microbe interactions, and the immunomodulatory functions of bacterial extracellular vesicles. AR is a member of the Marie Curie Alumni Association, International Society for Extracellular Vesicles (ISEV), the Bacterial EVs Task Force and Austrian Society for Extracellular Vesicles. IS - Associate Professor at the Institute of Specific Prophylaxis and Tropical Medicine, Medical University of Vienna. A parasitologist and immunologist by training, has extensive research experience in host–pathogen and host–microbiota interactions, with a focus on probiotics and extracellular vesicle biology. IS work combines immunology, microbiology, and nanobiotechnology to explore how microbial and parasite-derived molecules modulate immune responses. IS is a member of the International Society for Extracellular Vesicles (ISEV), the Austrian Society for Extracellular Vesicles, and the Austrian Society for Allergology and Immunology.

