## Supplement for "Extracellular vesicles as indicators of environmental stress response in *Lactiplantibacillus plantarum*: a multi-platform study"

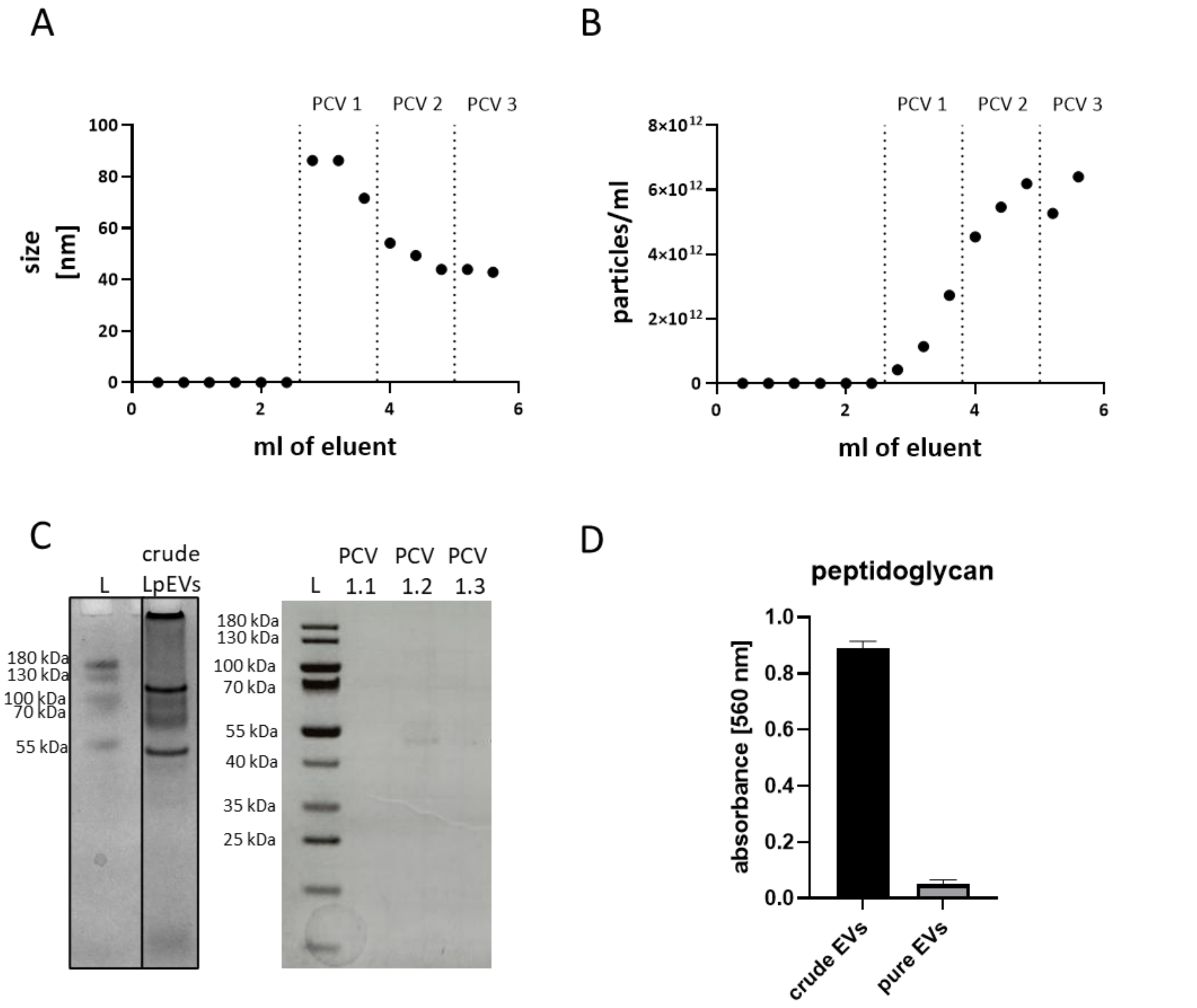


**Figure S1. Characteristics of LpEV fractions after SEC.** Each dot represents one fraction. (A) Representative chromatogram showing particle size in the isolated fractions. (B) Representative chromatogram showing particle concentration in the isolated fractions. PCV, purified collection volume. (C) SDS–PAGE analysis of crude LpEVs ($8 \times 10^9$ particles/well) and purified LpEVs (PCV 1.1 = $0.4 \times 10^9$ particles/well, PCV 1.2 = $7.6 \times 10^9$ particles/well, PCV 1.3 = $1.24 \times 10^9$ particles/well). (D) PGN measurement in $2.5 \times 10^{10}$ crude and purified LpEVs.

**Table S1. Batch-to-batch variability in the isolation and purification of LpEVs.** Data were collected from five independent batches, with bacterial cultures initiated from separate overnight starters.

| Batch number | Average particle size [nm] | Average particle concentration [particles/mL] |
|---|---|---|
| 1 | 69.11 | $1.5 \times 10^{12}$ |
| 2 | 83.74 | $2 \times 10^{12}$ |
| 3 | 78.09 | $0.7 \times 10^{12}$ |
| 4 | 66.8 | $3 \times 10^{12}$ |
| 5 | 71.76 | $1 \times 10^{12}$ |
| **Mean** | **73.9 ± 6.9** | **$1.64 \times 10^{12} \pm 0.91 \times 10^{12}$** |

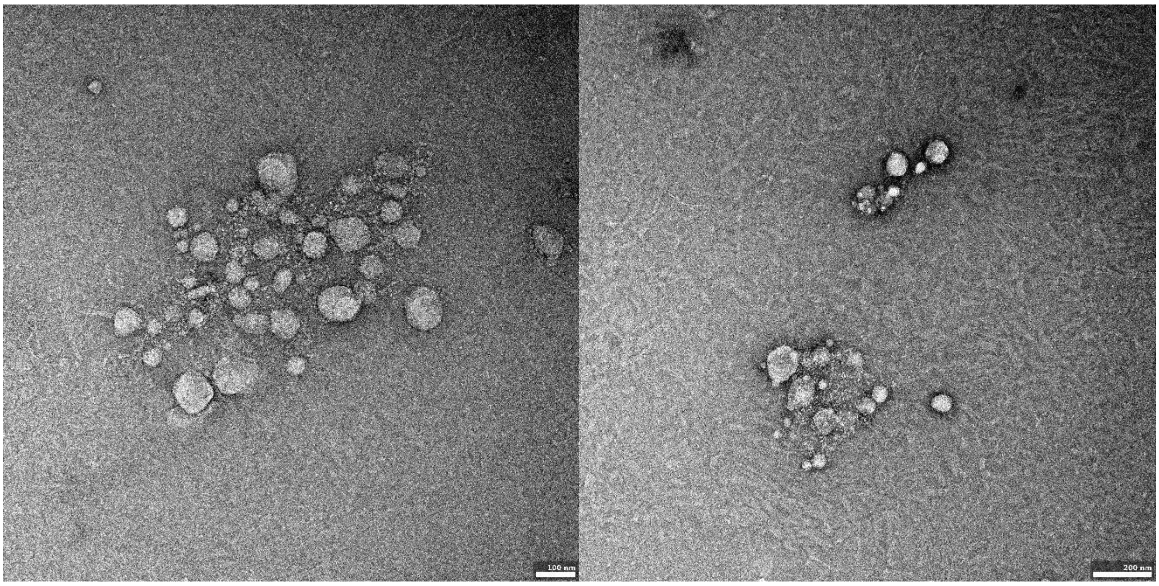


**Figure S2. TEM images of LpEVs shown at a wider field of view.**

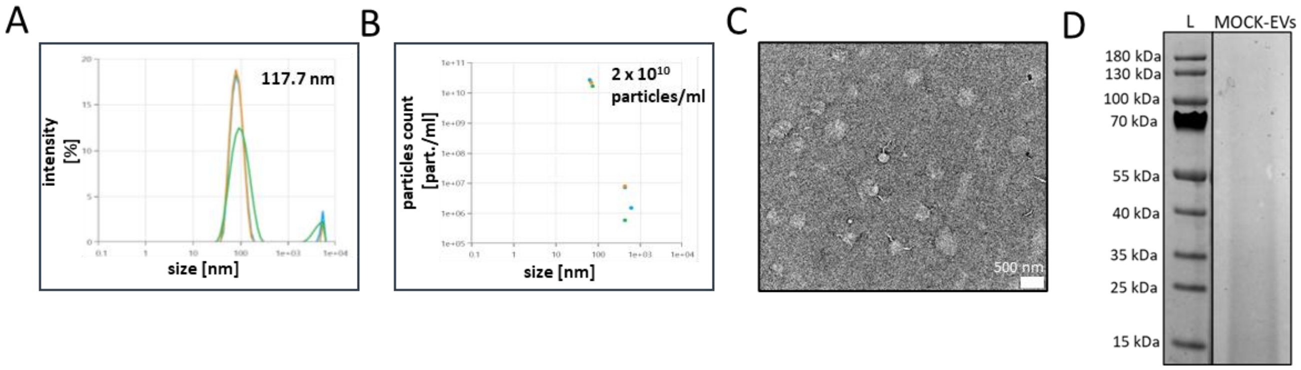


**Figure S3. Analysis of MOCK-EVs.** (A) Particle size distribution of MOCK-EVs measured using a Zetasizer. (B) Particle concentration of MOCK-EVs measured using a Zetasizer. (C) TEM images of MOCK-EVs. (D) SDS–PAGE analysis of a 20 µL MOCK-EV sample.

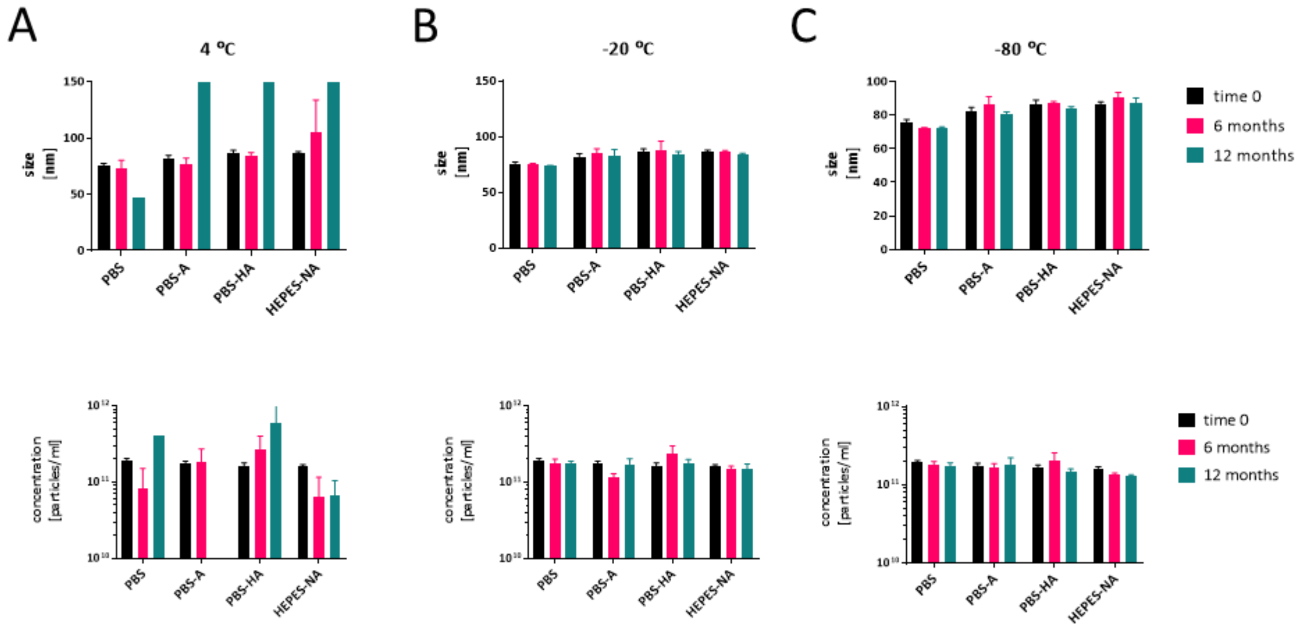


**Figure S4. Long-term stability of LpEVs under different buffer and temperature conditions.** (A) Particle size and concentration of LpEV samples stored at 4°C. (B) Buffer composition: PBS, phosphate-buffered saline; PBS-A, PBS with 0.2% BSA; PBS-HA, PBS-A with HEPES; HEPES-NA, HEPES with 0.9% NaCl and 0.2% BSA.

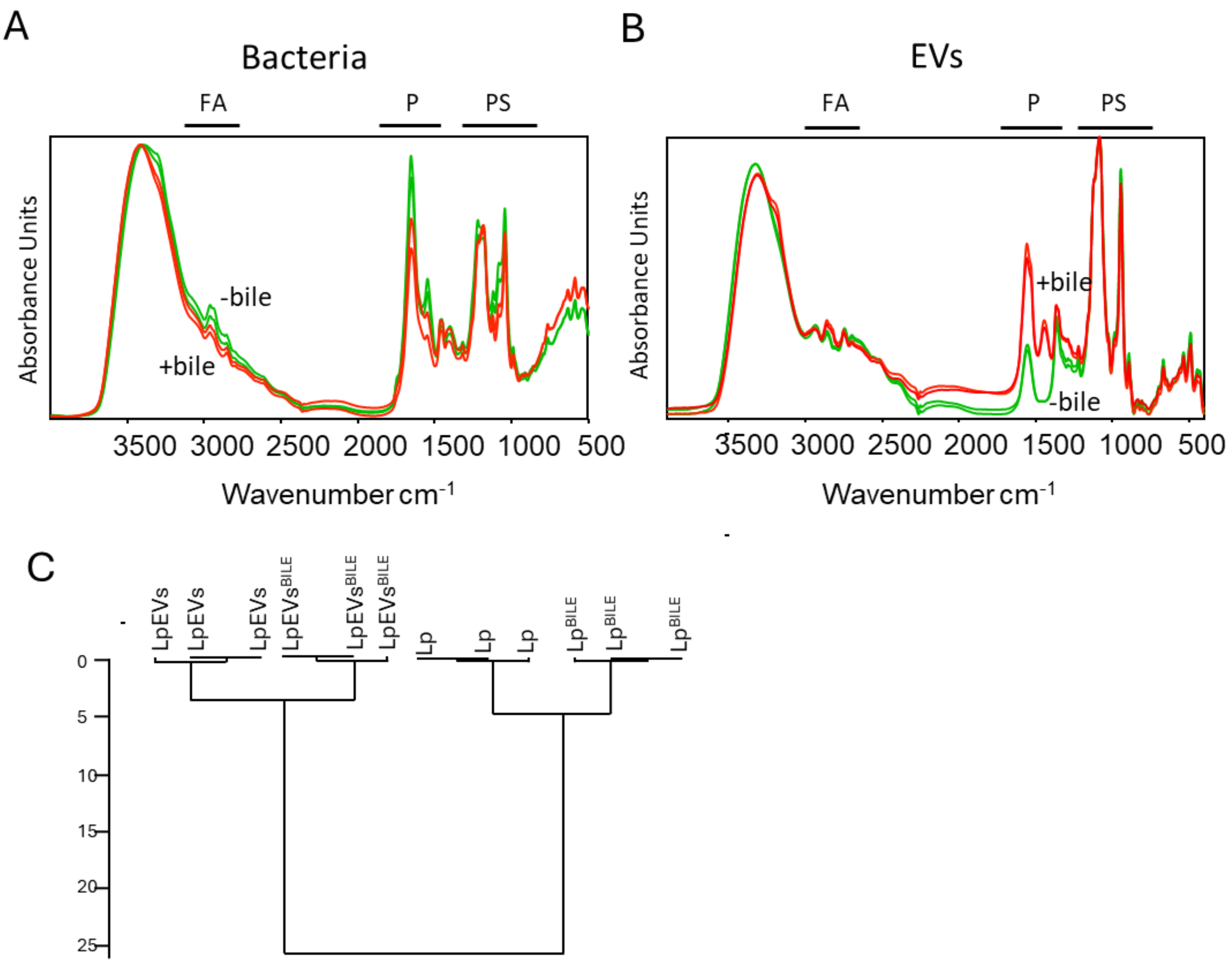


**Figure S5. Metabolic fingerprints of bacteria and EVs after cultivation in bile.** Baseline- and vector-normalized spectra from two additional biological replicates of Lp (green) and $Lp^{BILE}$ (red) (A) as well as LpEVs (green) and $LpEVs^{BILE}$ (red) (B) are shown. (C) HCA of the entire spectral range. FA, fatty acids; P, proteins; PS, polysaccharides.

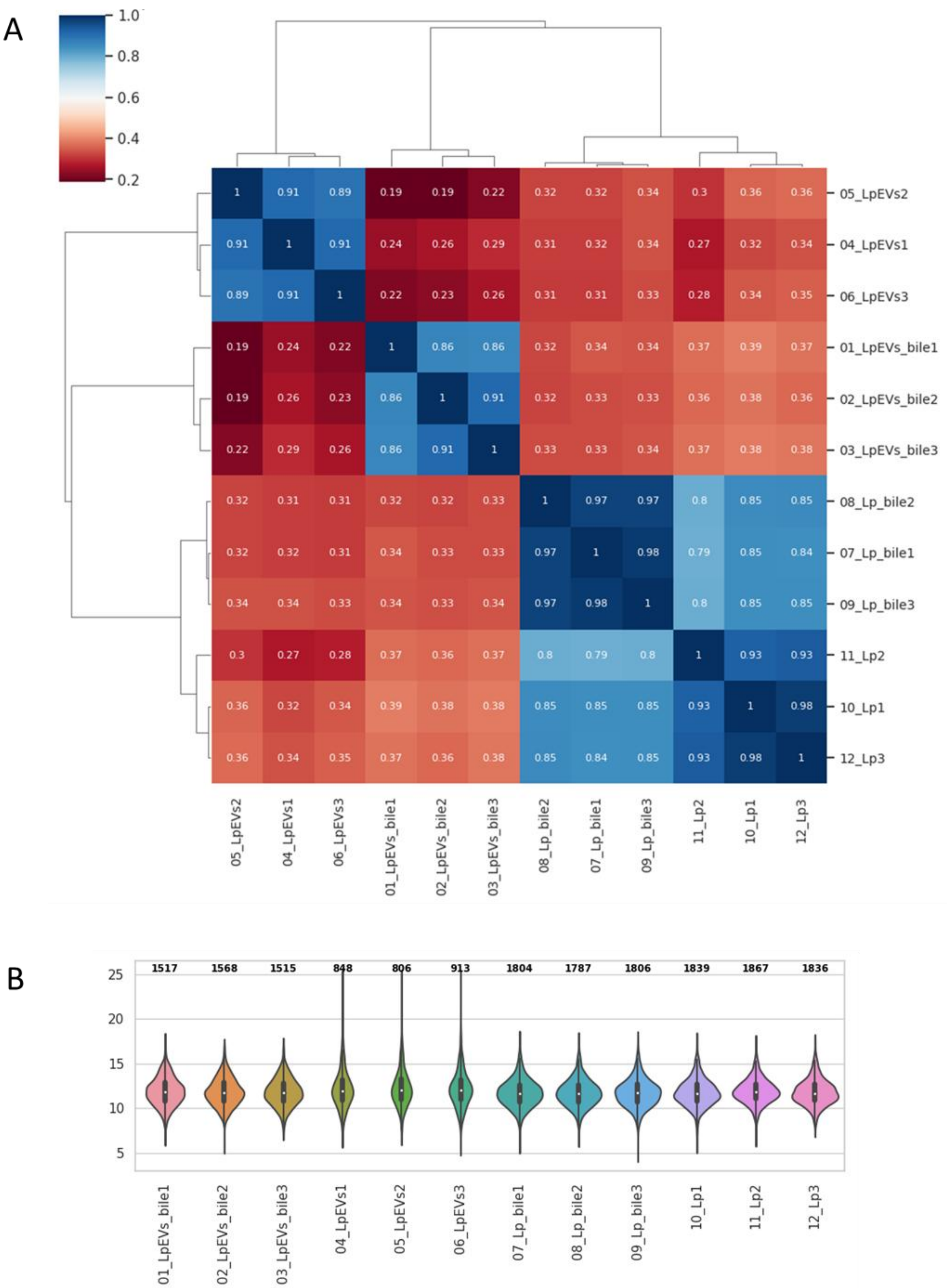


**Figure S6. Quality control data for proteomic analysis.** (A) Cluster map of pairwise Pearson correlation coefficients between biological and technical replicates (distance metric: Euclidean; clustering method: average linkage). (B) Violin plots showing the number of observed values.

**Table S2. STRING analysis of protein–protein interaction networks.**

| | Biological Process (Gene Ontology) | | | | |
|---|---|---|---|---|---|
| *GO-term* | *description* | *count in network* | *strength* | *signal* | *false discovery rate* |
| GO:0072524 | Pyridine containing compound metabolic process | 13 of 14 | 0.51 | 0.31 | 0.0469 |
| GO:0006400 | tRNA modification | 18 of 23 | 0.43 | 0.32 | 0.0360 |
| GO:0006364 | rRNA processing | 17 of 22 | 0.43 | 0.3 | 0.0449 |
| GO:0009451 | RNA modification | 30 of 42 | 0.39 | 0.41 | 0.0097 |
| GO:0034470 | ncRNA processing | 42 of 61 | 0.38 | 0.53 | 0.0013 |
| GO:0008033 | tRNA processing | 24 of 35 | 0.37 | 0.33 | 0.0280 |
| GO:0006396 | RNA processing | 43 of 65 | 0.36 | 0.5 | 0.0019 |
| GO:0034660 | ncRNA metabolic process | 57 of 89 | 0.34 | 0.59 | 0.00029 |
| GO:0006399 | tRNA metabolic process | 39 of 63 | 0.33 | 0.4 | 0.0097 |
| GO:0006767 | Water-soluble vitamin metabolic process | 26 of 42 | 0.33 | 0.3 | 0.0410 |
| GO:0016070 | RNA metabolic process | 71 of 128 | 0.28 | 0.52 | 0.00077 |
| GO:1901615 | Organic hydroxy compound metabolic process | 35 of 63 | 0.28 | 0.3 | 0.0360 |
| GO:0009165 | Nucleotide biosynthetic process | 47 of 88 | 0.27 | 0.34 | 0.0192 |
| GO:0019438 | Aromatic compound biosynthetic process | 87 of 178 | 0.23 | 0.44 | 0.0023 |
| GO:0034654 | Nucleobase-containing compound biosynthetic process | 67 of 137 | 0.23 | 0.35 | 0.0132 |
| GO:0006753 | Nucleoside phosphate metabolic process | 62 of 127 | 0.23 | 0.33 | 0.0190 |
| GO:1901362 | Organic cyclic compound biosynthetic process | 103 of 216 | 0.22 | 0.47 | 0.0012 |
| GO:0018130 | Heterocycle biosynthetic process | 96 of 198 | 0.22 | 0.47 | 0.0013 |
| GO:0055086 | Nucleobase-containing small molecule metabolic process | 71 of 149 | 0.22 | 0.35 | 0.0145 |
| GO:0009117 | Nucleotide metabolic process | 58 of 121 | 0.22 | 0.31 | 0.0258 |
| GO:0090407 | Organophosphate biosynthetic process | 58 of 125 | 0.2 | 0.28 | 0.0381 |
| GO:0046483 | Heterocycle metabolic process | 242 of 544 | 0.19 | 0.68 | 1.88e-07 |
| GO:0044281 | Small molecule metabolic process | 232 of 519 | 0.19 | 0.68 | 2.49e-07 |
| GO:0019637 | Organophosphate metabolic process | 85 of 188 | 0.19 | 0.34 | 0.0145 |
| GO:1901360 | Organic cyclic compound metabolic process | 250 of 568 | 0.18 | 0.67 | 1.88e-07 |
| GO:0006725 | Cellular aromatic compound metabolic process | 237 of 534 | 0.18 | 0.67 | 2.49e-07 |
| GO:0006139 | Nucleobase-containing compound metabolic process | 209 of 472 | 0.18 | 0.62 | 3.25e-06 |
| GO:0090304 | Nucleic acid metabolic process | 134 of 316 | 0.17 | 0.39 | 0.0044 |
| GO:0019752 | Carboxylic acid metabolic process | 102 of 240 | 0.17 | 0.32 | 0.0192 |
| GO:0044283 | Small molecule biosynthetic process | 97 of 228 | 0.17 | 0.31 | 0.0217 |
| GO:0044271 | Cellular nitrogen compound biosynthetic process | 125 of 301 | 0.16 | 0.34 | 0.0127 |
| GO:0006796 | Phosphate-containing compound metabolic process | 121 of 291 | 0.16 | 0.33 | 0.0142 |
| GO:0043436 | Oxoacid metabolic process | 103 of 243 | 0.16 | 0.32 | 0.0192 |
| GO:0006355 | Regulation of transcription, DNA-templated | 105 of 252 | 0.16 | 0.31 | 0.0227 |
| GO:0034641 | Cellular nitrogen compound metabolic process | 257 of 627 | 0.15 | 0.57 | 1.24e-05 |
| GO:0006793 | Phosphorus metabolic process | 125 of 306 | 0.15 | 0.32 | 0.0179 |
| GO:0019219 | Regulation of nucleobase-containing compound metabolic process | 106 of 257 | 0.15 | 0.3 | 0.0257 |
| GO:0050794 | Regulation of cellular process | 135 of 340 | 0.14 | 0.31 | 0.0217 |
| GO:0080090 | Regulation of primary metabolic process | 111 of 277 | 0.14 | 0.28 | 0.0360 |
| GO:0051171 | Regulation of nitrogen compound metabolic process | 111 of 276 | 0.14 | 0.28 | 0.0360 |
| GO:0031326 | Regulation of cellular biosynthetic process | 109 of 270 | 0.14 | 0.28 | 0.0360 |
| GO:0031323 | Regulation of cellular metabolic process | 111 of 276 | 0.14 | 0.28 | 0.0360 |
| GO:0010556 | Regulation of macromolecule biosynthetic process | 109 of 270 | 0.14 | 0.28 | 0.0360 |
| GO:0006082 | Organic acid metabolic process | 108 of 268 | 0.14 | 0.28 | 0.0360 |
| GO:0071704 | Organic substance metabolic process | 526 of 1352 | 0.13 | 0.73 | 1.16e-12 |
| GO:0044237 | Cellular metabolic process | 456 of 1179 | 0.13 | 0.67 | 2.82e-09 |
| GO:0065007 | Biological regulation | 159 of 410 | 0.13 | 0.31 | 0.0192 |
| GO:0019222 | Regulation of metabolic process | 113 of 286 | 0.13 | 0.27 | 0.0417 |
| GO:0060255 | Regulation of macromolecule metabolic process | 112 of 284 | 0.13 | 0.26 | 0.0449 |
| GO:0010468 | Regulation of gene expression | 110 of 279 | 0.13 | 0.26 | 0.0469 |
| GO:0008152 | Metabolic process | 584 of 1512 | 0.12 | 0.76 | 5.17e-15 |
| GO:0044238 | Primary metabolic process | 433 of 1128 | 0.12 | 0.64 | 4.16e-08 |
| GO:0050789 | Regulation of biological process | 149 of 387 | 0.12 | 0.29 | 0.0267 |
| GO:0006807 | Nitrogen compound metabolic process | 350 of 930 | 0.11 | 0.52 | 3.34e-05 |
| GO:0009058 | Biosynthetic process | 233 of 627 | 0.11 | 0.34 | 0.0097 |
| GO:0044249 | Cellular biosynthetic process | 217 of 579 | 0.11 | 0.34 | 0.0102 |
| GO:1901576 | Organic substance biosynthetic process | 221 of 607 | 0.1 | 0.3 | 0.0217 |
| GO:1901564 | Organonitrogen compound metabolic process | 215 of 600 | 0.09 | 0.27 | 0.0360 |

| | Cellular Component (Gene Ontology) | | | | |
|---|---|---|---|---|---|
| *GO-term* | *description* | *count in network* | *strength* | *signal* | *false discovery rate* |
| GO:0005829 | Cytosol | 147 of 299 | 0.23 | 0.68 | 1.65e-06 |
| GO:0005737 | Cytoplasm | 519 of 1137 | 0.2 | 0.96 | 3.29e-27 |
| GO:0005622 | Intracellular anatomical structure | 543 of 1232 | 0.18 | 0.93 | 2.86e-26 |
| GO:0110165 | Cellular anatomical entity | 741 of 2390 | 0.03 | 0.32 | 0.0092 |

| | Local Network Cluster (STRING) | | | | |
|---|---|---|---|---|---|
| *cluster* | *description* | *count in network* | *strength* | *signal* | *false discovery rate* |
| CL:453 | Mixed, incl. Catalytic activity, acting on a nucleic acid, and DNA repl... | 58 of 103 | 0.29 | 0.31 | 0.0332 |

| | KEGG Pathways | | | | |
|---|---|---|---|---|---|
| *pathway* | *description* | *count in network* | *strength* | *signal* | *false discovery rate* |
| lpl01100 | Metabolic pathways | 209 of 510 | 0.15 | 0.48 | 0.00031 |

| | Subcellular Localization (COMPARTMENTS) | | | | |
|---|---|---|---|---|---|
| *compartment* | *description* | *count in network* | *strength* | *signal* | *false discovery rate* |
| GOCC:0043231 | Intracellular membrane-bounded organelle | 58 of 116 | 0.24 | 0.3 | 0.0320 |
| GOCC:0005829 | Cytosol | 189 of 416 | 0.2 | 0.63 | 3.00e-06 |
| GOCC:0005737 | Cytoplasm | 382 of 860 | 0.19 | 0.84 | 1.22e-14 |
| GOCC:0005622 | Intracellular | 508 of 1235 | 0.15 | 0.81 | 1.63e-16 |

| | Annotated Keywords (UniProt) | | | | |
|---|---|---|---|---|---|
| *keyword* | *description* | *count in network* | *strength* | *signal* | *false discovery rate* |
| KW-0285 | Flavoprotein | 24 of 30 | 0.44 | 0.42 | 0.0098 |
| KW-0819 | tRNA processing | 18 of 23 | 0.43 | 0.31 | 0.0435 |
| KW-0436 | Ligase | 50 of 82 | 0.32 | 0.43 | 0.0052 |
| KW-0560 | Oxidoreductase | 89 of 181 | 0.23 | 0.41 | 0.0052 |
| KW-0804 | Transcription | 64 of 142 | 0.19 | 0.27 | 0.0458 |
| KW-0963 | Cytoplasm | 115 of 262 | 0.18 | 0.39 | 0.0052 |
| KW-0238 | DNA-binding | 82 of 185 | 0.18 | 0.31 | 0.0250 |
| KW-0378 | Hydrolase | 146 of 350 | 0.16 | 0.38 | 0.0052 |
| KW-0808 | Transferase | 177 of 478 | 0.11 | 0.26 | 0.0435 |

*(less ...)*
