## Supplementary material for "Extracellular vesicles as indicators of environmental stress response in *Lactiplantibacillus plantarum*: a multi-platform study": Information on the access to databases

Information on the access to the databases

EV-TRACK

You may access and check the submission of experimental parameters to the EV-TRACK knowledgebase via the following URL: <http://evtrack.org/review.php>. Please use the EV-TRACK ID (EV250022) and the last name of the first author (Razim) to access our submission.

Proteomics

Log in to the PRIDE website using the following details:

**Project accession:** PXD060959

**Token:** We15zOWWdqms

Alternatively, reviewer can access the dataset by logging in to the PRIDE website using the following account details:

**Username:** 

**Password:** 2hbqc96iuR49
